# Understanding the impact of third-party species on pairwise coexistence

**DOI:** 10.1101/2022.07.12.499717

**Authors:** Jie Deng, Washington Taylor, Serguei Saavedra

**Author notes:** **Author contributions:** J.D. and S.S designed research; J.D was responsible for investigation; all authors were responsible for conceptualization, formal analysis, methodology, software, validation, writing, reviewing, and editing; S.S was responsible for funding acquisition and supervision. **Competing financial interests** The authors declare no competing financial interests. The funders had no role in study design, data collection and analysis, decision to publish, or preparation of the manuscript.

## Abstract

The persistence of virtually every single species depends on both the presence of other species and the specific environmental conditions in a given location. Because in natural settings many of these conditions are unknown, research has been centered on finding the fraction of possible conditions (probability) leading to species coexistence. The focus has been on the persistence probability of an entire multispecies community (formed of either two or more species). However, the methodological and philosophical question has always been whether we can observe the entire community and, if not, what the conditions are under which an observed subset of the community can persist as part of a larger multispecies system. Here, we derive long-term (using analytical calculations) and short-term (using simulations and experimental data) system-level indicators of the effect of third-party species on the coexistence probability of a pair (or subset) of species under unknown environmental conditions. We demonstrate that the fraction of conditions incompatible with the possible coexistence of a pair of species tends to become vanishingly small within systems of increasing numbers of species. Yet, the probability of pairwise coexistence in isolation remains approximately the expected probability of pairwise coexistence in more diverse assemblages. In addition, we found that when third-party species tend to reduce (resp. increase) the coexistence probability of a pair, they tend to exhibit slower (resp. faster) rates of competitive exclusion. Long-term and short-term effects of the remaining third-party species on all possible specific pairs in a system are not equally distributed, but these differences can be mapped and anticipated under environmental uncertainty.

**Author Summary:** It is debated whether the frequency with which two species coexist in isolation or within a single environmental context is representative of their coexistence expectation within larger multispecies systems and across different environmental conditions. Here, using analytical calculations, simulations, and experimental data, we show why and how third-party species can provide the opportunity for pairwise coexistence regardless of whether a pair of species can coexist in isolation across different environmental conditions. However, we show that this opportunity is not homogeneously granted across all pairs within the same system. We provide a framework to understand and map the long-term and short-term effects that third-party species have on the coexistence of each possible subset in a multispecies system.

## Introduction

The persistence of virtually all living organisms on Earth depends to a greater or lesser extent on the presence of other living organisms and on the environmental (abiotic and biotic) conditions present in a given place and time (1, 2). This observation has established a rich research program in quantifying pairwise interactions and their impact on pairwise coexistence across environmental gradients (3–8). In this line, observational work has been focused on finding the frequency of occurrence of multiple pairs of species across different environments either in isolation or within different systems of multiple species (9–16). For example, studies have long debated whether there exists in nature pairs of species whose niches forbid their coexistence regardless of the environment—known as the checkerboard hypothesis (11, 17). Instead, experimental studies have shown that pairwise coexistence strongly depends on the details of both the system and the environment (10, 12, 18, 19). In fact, it has been shown that the coexistence expectations that may operate under controlled conditions (or unique environments) (12, 14) do not necessarily operate under uncontrolled conditions (or diverse environments) (5, 20). These observational results have shown that in order to implement successful interventions in ecological communities, it is essential to develop a testable theory to be able to explain why and how emergent processes at the system level can affect the possibility that a given pair (or community) of species will coexist under different abiotic and biotic environments.

Because the environmental conditions acting on species are typically unknown and diverse in natural settings, the majority of theoretical work has been centered on deriving the fraction of conditions (set of parameter values) leading to the persistence of a community (formed by either two or more species) (8, 21–23). Yet, species are seldom in isolation and the coexistence of a specific combination of multiple species is expected to be rare in a random environment (5, 24, 25). Moreover, the methodological and philosophical question has always been whether we can observe the entire community and, if not, what the conditions are under which an observed subset of the community can persist as part of a larger multispecies system. The answer to this question, however, may also be context-dependent due to the presence and absence of third-party species, which can be acting as ecosystem engineers (26, 27). Thus, it is unclear how far knowledge of pairwise coexistence in isolation (or full community persistence) can take us while investigating ecological systems, and how we should compare the effects of third-party species on the coexistence probability of a pair (or community) of species.

In this line, ecological research has suggested that species embedded into larger multispecies systems (more than two species) may experience higher-order effects, i.e., the effect of species *i* on the per capita growth rate of species *j* might itself depend on the abundance of a third species *k* due to either compensatory effects, supra-additivity, trait-mediated effects, functional effects, meta-community effects, or indirect effects (8, 28–31). Unfortunately, additional research has shown that it is virtually impossible to derive those potential effects since many parameter combinations can equally explain the observed ecological dynamics—a problem known as structural identifiability (32, 33). In fact, many of the interactions measured under multispecies systems are not direct effects, as studies often believe (15, 16), but chains of direct effects (34, 35). Moreover, measurements of species interactions are expected to change as a function of the type of data (i.e., snapshot or time), experiment (e.g., long term or short term), perturbations (e.g., pulse or press), and dimensionality, leading to inconsistencies and inference issues (32, 36, 37). Thus, it is unclear whether the frequency with which two or more species coexist in isolation is representative of their coexistence expectation within larger multispecies systems and across different environments.

Here, we follow a geometric and probabilistic analysis based on nonlinear dynamics at equilibrium to estimate the effect of third-party species (the larger multispecies system) on the coexistence of a pair (community) of species under unknown environmental conditions using only information from pairwise species interactions. We estimate these system-level effects over the long term (using analytical calculations) and over the short term (using simulations and available experimental data on gut microbiota). Building on generalized Lotka-Volterra (gLV) dynamics (8), we provide a geometric understanding regarding whether the conditions limiting the coexistence of a pair of species in isolation (the species pool consists only of this pair) remain with third-party species (when the species pool increases). Next, we derive a system-level indicator of the extent to which third-party species can affect the probability of coexistence of a pair relative to its probability in isolation. Then, we compare the numerical and experimental effects to the analytical expectation in order to distinguish the role that third-party species play in shaping pairwise coexistence under short-term and long-term behavior, respectively. Finally, we show how our theory can be used to provide a cartographic representation (38) of the long-term and short-term effects acting on each pair within a multispecies system under environmental uncertainty. While our focus is on pairwise coexistence (following traditional work in ecology), our methodology and results are scalable to any subset (community) dimension.

## Methods

### Traditional indicators of pairwise coexistence

Our theoretical framework is based on the tractability of the gLV dynamics (33), where the percapita growth rate of species *i* ∈ {1, ⋯, |𝒮|} ≡ 𝒮 is written as *f*_*i*_(**N**; ***θ***) = *θ*_*i*_ − Σ_*j*∈𝒮_ *a*_*ij*_*N*_*j*_, where the vector **N** = (*N*_1_, …, *N*_|𝒮|_)^⊺^ represents the density of all species in the system 𝒮. The matrix **A** = (*a*_*ij*_) ∈ ℝ^|𝒮|×|𝒮|^ corresponds to the interaction matrix dictating the internal structure of the system, i.e., the per-capita effect of species *j* on an individual of species *i*. The effective parameters ***θ*** = (*θ*_1_, …, *θ*_|𝒮|_)^⊺^ ∈ ℝ^|𝒮|^ consist of the phenomenological effect of the internal (e.g., intrinsic growth rate), abiotic (e.g., temperature, pH, nutrients), and biotic factors (e.g., unknown species) acting on the per-capita growth rate of each particular species. That is, we consider that the *effective growth rate* ***θ*** represents the total additional effect of all the unknown potential factors as a function of the environmental conditions acting on each species independently. Note that pairwise effects *a*_*ij*_ from a given pool of species are assumed to be known. Our essential assumptions are thus that the effective growth rates ***θ*** are environmentally determined (i.e., change with the environment), but a pairwise interaction *a*_*ij*_ is a property only of the species pair (i.e., invariant under change of environment or addition of further species).

Under steady-state dynamics (i.e., ***f*** = 0), the necessary (but not sufficient) condition for species coexistence is the existence of a feasible equilibrium (i.e., ***N***^*^ = **A**^−1^***θ*** *>* 0) (22). We are interested here in globally stable systems; feasible and globally stable systems fulfill the necessary and sufficient conditions for coexistence (22). The combinations of the effective growth rates ***θ*** compatible with the feasibility of a multispecies system S (known as the feasibility region *D*_*F*_) can be described as:

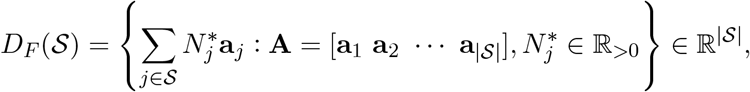

where **a**_*j*_ is the *j*^th^ column vector of the interaction matrix **A**. Additionally, assuming no a priori information about how the environmental conditions affect the effective growth rate (39), the feasibility region can be normalized by the volume of the entire parameter space (25) as

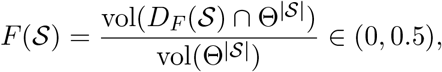

where Θ^|𝒮|^ is the (|𝒮| − 1)-dimensional closed unit sphere in dimension |𝒮|. Geometrically, the feasibility region is the convex hull of the |𝒮| spanning vectors in |𝒮|-dimensional space, and thus always corresponds to an |𝒮|-dimensional solid angle that is contained within a single hemisphere (e.g., an acute angle in the case |𝒮| = 2). Thus, the feasibility that is normalized by the entire |𝒮|-dimensional space cannot be higher than 0.5. Ecologically, this normalized feasibility region can be interpreted as the probability of feasibility of a system 𝒮 characterized by known pairwise interactions **A** under environmental uncertainty (all possible effective growth rates are equally likely to happen). The measure *F* (𝒮) can be efficiently estimated analytically (25, 40) or using Monte Carlo methods (Sec. S4). Moreover, *F* (𝒮) is robust to gLV dynamics with linear functional responses (25), a large family of nonlinear functional responses (39, 41), gLV stochastic dynamics (39, 41), and as a lower bound for complex polynomial models in species abundances (33).

*F* (𝒮) represents the feasibility of all |𝒮| species together. That is, if a system (i.e., a given pool) is composed of 2 species or of 10 species, *F* (𝒮) represents the probability of feasibility of the full set of 2 or 10 species, respectively (25). In the case where |𝒮| = 2, *F* (𝒮) corresponds to the traditional indicator of pairwise coexistence in isolation, which is proportional to what is typically known as niche overlap in gLV competition systems (42, 43). It is known that *F* (𝒮) will tend to decrease as a function of the dimension of the system (23, 33): *F* (𝒮_1_) *> F* (𝒮_2_) if |𝒮_1_| < |𝒮_2_|. Thus, focusing on a given pair of species *Ƶ* = {*i, j*} within a system *Ƶ* ⊂ 𝒮, *F* (𝒮) will likely underestimate the probability of feasibility for the pair, whereas using only the feasibility of the pair in isolation *F* (*Ƶ*) can be potentially misleading. For example, considering a pair that has a rather small feasibility region in isolation (*F* (*Ƶ*) ∼ 0, e.g., almost identical niches) will prompt us to conclude that the coexistence of this pair should be forbidden across all possible environmental conditions—as suggested by the checkerboard hypothesis (11, 14, 17). But, can this pair of species coexist more easily with additional third-party species (44, 45)? And will different pairs within the same system of multiple species be equally benefited (or affected)?

### Looking at traditional indicators from a systems perspective

To understand the extent to which *F* (*Ƶ*) provides a reliable indicator of the possibility of coexistence of a pair (or community) *Ƶ* = {*i, j*} within a larger multispecies system 𝒮 characterized by a pairwise interaction matrix **A**, we study geometrically how the feasibility conditions ***θ*** change when a pair of species is in isolation and with third-party species. We approach this question from several different perspectives. First, we start by asking to what extent can third-party species act as ecosystem engineers and modify the environment into a more suitable habitat (26, 27)? In particular, for a given pair *i, j* within a fixed community containing third-party species, we consider the range of environmental conditions associated with the two parameters *θ*_*i*_, *θ*_*j*_ such that it is *possible* for the pair *i, j* to exist for *some* set of values for the remaining *θ*_*k*_’s. This gives a sense of the potential environmental range of this pair in the context of a fixed community. In the following subsection we explore the more precise question of what the *probability* of coexistence is when we average over all possible values of all the *θ*_*k*_’s.

Mathematically, the projection region *D*_Proj_ of the feasibility region *D*_*F*_ of a multispecies system 𝒮 onto the 2-dimensional parameter space of a pair of species *Ƶ* can be represented by the conical hull, i.e., the set of all conical combinations of the projection of spanning rays:

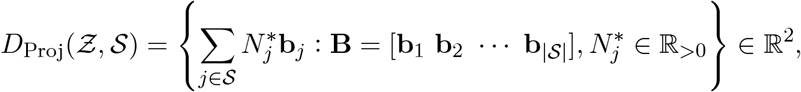

where **b**_*j*_ is the *j*^th^ column vector of **B**, and it contains a subset of elements corresponding to the pair *Ƶ* in the column vector **a**_*j*_ of the interaction matrix **A**. In other words, **B** consists of the row vectors **a**_*i*_ of **A** that correspond to the pair *Ƶ* (i.e., *i* ∈ *Ƶ*). For example, for the pair of species 1 and 3 (i.e., *Ƶ* = {1, 3}), for each column vector **a**_*j*_ = (*a*_1*j*_, ⋯, *a*_|𝒮|*j*_)^⊺^, *j* ∈ {1, ⋯, |𝒮|} in **A**, we have **b**_*j*_ = (*a*_1*j*_, *a*_3*j*_)^⊺^ in **B**. Then, we can obtain the normalized projection by the following ratio of volumes:

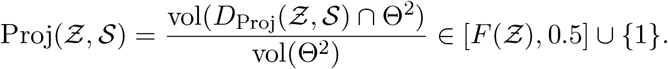

Recall that *F* (*Ƶ*) is the feasibility of pair *Ƶ* in isolation (using the corresponding 2-dimensional submatrix **A**[*i, j*; *i, j*]). This projection region captures the first ecological question raised above: what is the range of *θ*_*i*_, *θ*_*j*_ under which it is *possible* for species *i, j* to coexist for a given matrix **A** and some values of the remaining parameters *θ*_*k*_, *k*≠ *i, j*.

If the projection region is not the entire 2-dimensional parameter space ***θ*** of the pair, the projection ranges within Proj ∈ (0, 0.5] (Fig. 1A provides an illustration). Otherwise, Proj = 1, indicating that the region (*θ*_1_, *θ*_2_) incompatible with the possible coexistence of the pair disappears. In other words, in these cases pairwise coexistence can be possible under any combination of (*θ*_*i*_, *θ*_*j*_). The realization of such coexistence, however, still depends on the other ***θ*** values associated with the third-party species in the system. In fact, per definition *D*_Proj_(*Ƶ*, 𝒮) ≥ *D*_*F*_ (*Ƶ*), the entire projection region *D*_Proj_(*Ƶ*, 𝒮) defines equal or weaker constraints on the plane (*θ*_*i*_, *θ*_*j*_) for the coexistence of the pair. Thus, the amount of *projection contribution* increasing the feasibility region on the plane (*θ*_*i*_, *θ*_*j*_) can be defined as

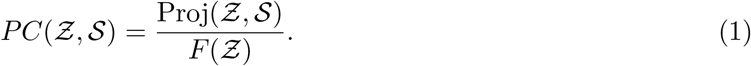

**Figure 1:**
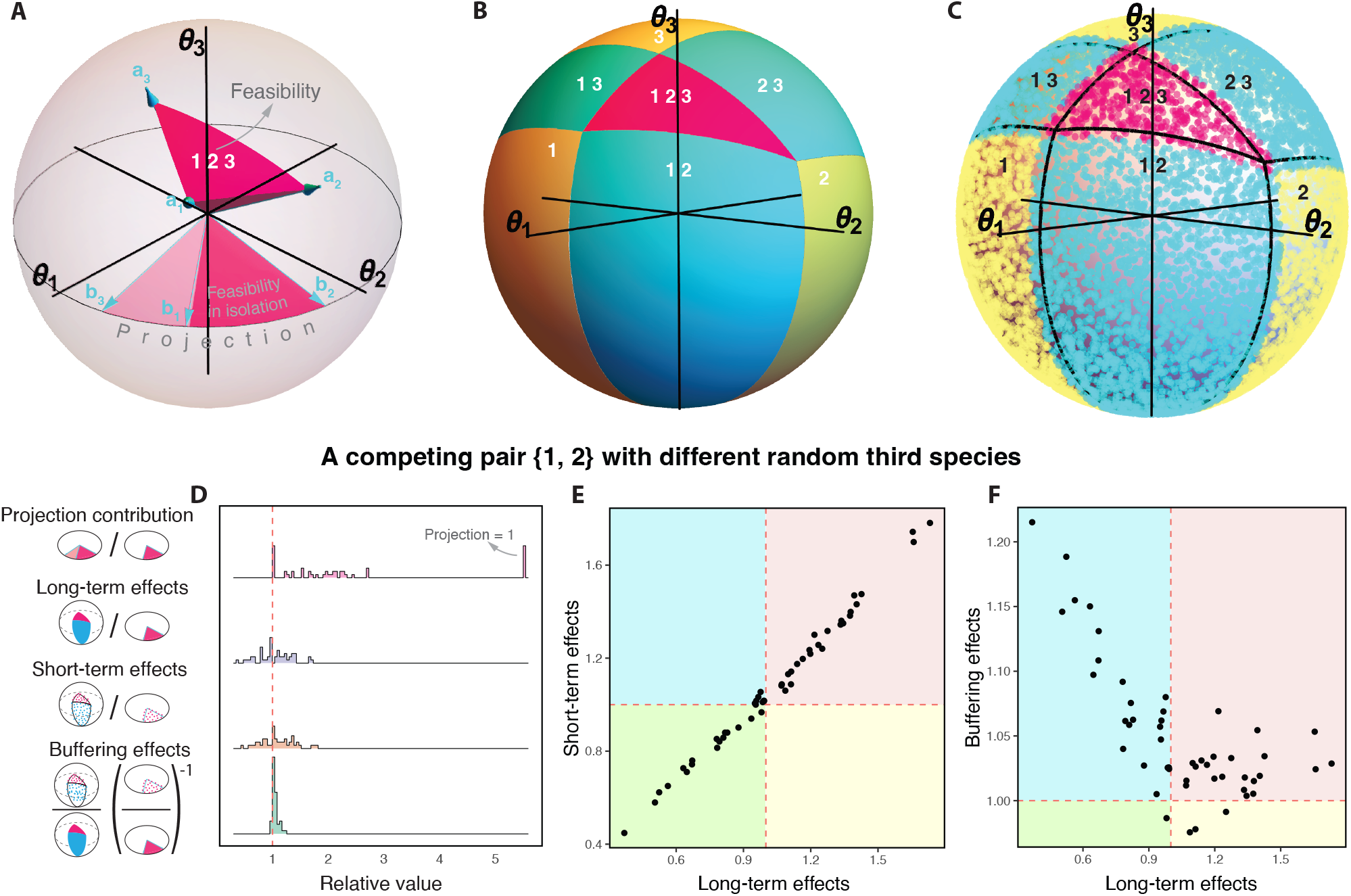
Geometric perspective of the effect of third-party species on pairwise coexistence. For a fictitious system of three species, Panel **A** shows the geometrical representation of the feasibility region *D*_*F*_ (𝒮) (i.e., pink cone spanned by **a**_1_, **a**_2_, **a**_3_) of an illustrative three-species system characterized by the interaction matrix **A** ℝ^3×3^ in the three-dimensional space of effective growth rates (*θ*_1_, *θ*_2_, *θ*_3_). The panel also shows the projection *D*_Proj_(*Ƶ*, 𝒮) of the spanning rays onto the two-dimensional plane (*θ*_1_, *θ*_2_) of the pair *Ƶ* = {1, 2} (i.e., light pink and dark pink disk spanned by **b**_2_, **b**_3_). The feasibility of pair {1, 2} in isolation (i.e., a 2-species pool) *D*_*F*_ (*Ƶ*) is given by **b**_1_, **b**_2_ (i.e., dark pink disk), and becomes a fraction of the projection corresponding to the possible coexistence of species 1 and 2 only. Panel **B** illustrates the geometric partition of the analytical feasibility regions of all possible compositions in the previous three-species system. For the same system, the points in Panel **C** show the resulting species composition (species with final abundances *N* < 10^−6^ are considered statistically extinct) from 10,000 simulations over a short finite time interval (total time = 200, step size = 0.01) conducted by the Runge-Kutta method. Note that these results are a function of the time span of simulations (see Fig. S7), although the qualitative effects are similar for other choices of time span. The black lines correspond to the borders of the analytical feasibility regions in Panel **B**. Panel **D** shows the distribution of system-level effects calculated on a fixed hypothetical pair of symmetrically competing species *Ƶ* = {1, 2} when assembled with different random third species. The distributions are generated using 50 random third species (see text for details). Each point in the distributions corresponds to the same pair with a different set of third-party species. For reference, the dashed line shows the value of one: the relative feasibility in isolation. Panels **E-F** show the positive and negative association of the long-term effect with the short-term and buffering effects, respectively (Panel **E**: Pearson’s product-moment correlation = 0.995, *p*-value < 0.001; Panel **F**: Pearson’s product-moment correlation = −0.707, *p*-value < 0.001). These results are complemented by a perturbative analysis for weakly interacting systems in Sec. S5.

Because the combination of spanning vectors increases as the dimension of the system |𝒮| grows, the projection Proj(*Ƶ*, 𝒮) is expected to approach 1 in large multispecies systems (Figs. S3-S4).

### Analytical system-level effects

While the projection contribution, which measures the *possibility* of pairwise coexistence, is always beneficial to the likelihood of pairwise coexistence, it is measured in the two-dimensional (*θ*_*i*_, *θ*_*j*_) plane of the pair {*i, j*} in question. The actual probability of pairwise coexistence, however, depends on the ***θ*** values of the rest of the species in the community. To find long-term analytical indicators of the effect of third-party species on pairwise coexistence, first we propose to do a geometric partition of the parameter space of ***θ*** (a closed unit sphere) into non-invadable feasibility regions of all possible species compositions within a multispecies system. These regions are disjoint when there is a global stable equilibrium. Formally, in a multispecies system 𝒮 with an interaction matrix **A** and a stable global equilibrium, the feasibility region of a composition 𝒞 (a subset of species found in the system, i.e., 𝒞 ⊆ 𝒮, 𝒞 ≠ ∅) can be defined as:

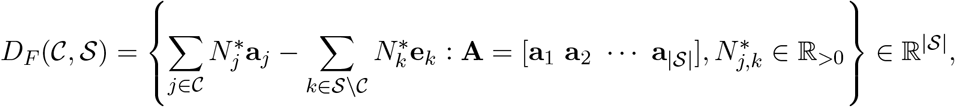

where **e**_*k*_ is a unit vector whose *k*^th^ entry is 1 and 0 elsewhere. In fact, for a composition 𝒞, we only need the subset of column vectors **a**_*j*_ of **A** corresponding to the composition 𝒞 (i.e., *j* ∈ 𝒞) instead of the whole matrix. For example, for the species composition 𝒞 = {1, 2, 3}, only the 1^st^, 2^nd^ and 3^rd^ column vectors of **A** (i.e., the sub-matrix [**a**_1_ **a**_2_ **a**_3_]) are needed to define its feasibility region. Furthermore, we can define the normalized feasibility of a composition 𝒞 by

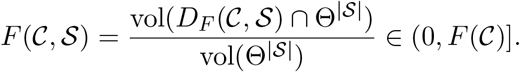

*F* (𝒞) is a function of the sub-matrix formed only by the interactions between the species in C (i.e., **A**[𝒞; 𝒞]); while *F* (𝒞, 𝒮) is a function of this sub-matrix augmented by the system-level effects of the composition on the third-party species in the system. In general, *F* (𝒞, 𝒮) = *F* (𝒞) only when 𝒞 = 𝒮. Note that when the full community 𝒮 admits a globally stable equilibrium, non-invadable equilibria of any subsystems 𝒞 are also stable (46).

Following our geometric partition, we define the probability of feasibility of a pair of species *Ƶ* in a larger multispecies system 𝒮 as the sum of all the feasibility regions where the pair forms part of the species composition (*Ƶ* ⊆ 𝒞 ⊆ 𝒮):

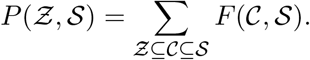

Because the proposed probability of feasibility is related to the solution of the system at equilibrium, we estimate analytically the *long-term effect* of a multispecies system 𝒮 on a pair *Ƶ* by

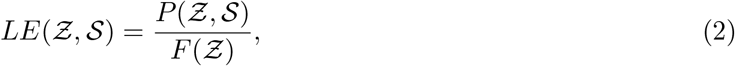

which represents the extent to which third-party species can modify the probability of pairwise coexistence relative to the probability of the pair in isolation allowing infinite time (Fig. 1B). Note that the long-term effect given by Eq. (2) describes the effect of the larger multispecies system on the *probability* that the composition 𝒞 arises as a community of the multispecies system, while the projection contribution given by Eq. (1) denotes the effect of the larger multispecies system in weakening constraints on the range of parameters allowing *possible* coexistence of composition 𝒞. The actual probability of pairwise coexistence depends on averaging over the effective growth rates ***θ*** of the rest of the species in the community. The long-term effect of Eq. (2) takes that into account and therefore, measures the relative probability of a pairwise equilibrium being feasible within a specific multispecies community. Note also that while the projection contribution always expands the possible range of coexistence for a given pair, the long-term effect that measures the average coexistence range is often (roughly half of all cases) decreased; these statements are completely consistent since the probability of the additional *θ*_*k*_’s taking values that significantly expand the pairwise coexistence range can be nonzero but quite small compared to the probability of contracting the coexistence range, and the projection contribution only incorporates the former.

### Numerical system-level effects

While the long-term behavior of a multispecies system provides an expectation of the effect of thirdparty species on a pair of species under an ideal setting; in practice, the study of pairwise coexistence in natural or experimental settings depends on the time scale of investigation, extinction thresholds, and the finite number of replicates. This implies that transient dynamics may play an additional role and observed coexistence rates may be different from the long-term expectations (12, 47). Thus, to provide a practical short-term perspective regarding the effect of third-party species on the probability of pairwise coexistence, we complement the analytical calculations of feasibility with numerical calculations. We simulate gLV dynamics over a given time interval across an arbitrary number of repetitions with arbitrary initial conditions (picking from a uniform distribution between 0 and 1 for each species, but starting with the same abundance for all species yields similar results) and classify species with a final abundance *N* < 10^−6^ (results are robust to different thresholds) as statistically extinct (Fig. 1C). The time interval 200 is chosen for consistency with the experimental data in Sec. S1. That is, while correlations between short-term and experimental effects were in general good, correlations using a total time of 200 were the strongest (Figs. S10-S11), given the specific time parameters of the experiments. Hence, for consistency, we simply set all time intervals of simulations to 200. For multispecies systems, the vector of effective growth rates ***θ*** is sampled randomly from the closed unit sphere. Then, to control for confounding factors, we take the two values in the sampled vector ***θ*** that correspond to the pair as the effective growth rates for the pair in isolation. All simulations are conducted by the Runge-Kutta method.

We quantify the *short-term effect* of a multispecies system 𝒮 on a pair of species *Ƶ* by the ratio of the simulated frequencies of pairwise coexistence with third-party species and the simulated frequencies in isolation:

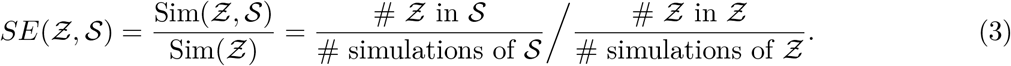

The simulated coexistence frequencies are always greater than or equal to the analytical, i.e., there is a short-term buffering effect Sim(*Ƶ*, 𝒮) ≥ *P* (*Ƶ*, 𝒮) and at Sim(*Ƶ*) ≥ *F* (*Ƶ*), because we always start with all species present in the system and the global equilibrium is assumed to be stable. Only after a sufficiently long time do the simulated and analytical frequencies become the same (Fig. S7).

To study whether the equilibration between short-term and long-term effects happens more quickly with third-party species or in isolation, we quantify the *buffering effect* of a simulated system by the ratio between the short-term and long-term effects

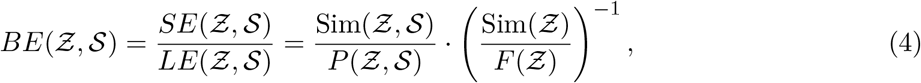

which represents the transient benefit (if *>* 1) or disadvantage (if < 1) to a pair of species after a finite time simulation of the system dynamics relative to the expected long-term behavior in the analytical calculations. For a given pair, the value of the buffering effect can change according to the chosen criteria for the simulations (Fig. S10), particularly the time span of the simulation. Yet, *BE*(*Ƶ*, 𝒮) = 1 if the two simulated frequencies (within systems Sim(*Ƶ*, 𝒮) and in isolation Sim(*Ƶ*)) are greater than their analytical counterparts (within systems *P* (*Ƶ*, 𝒮) and in isolation *F* (*Ƶ*)) in equal proportion. On the other hand, if *BE*(*Ƶ*, 𝒮) is greater (resp. less) than 1, the buffering effect is stronger with third-party species (resp. in isolation).

### Comparing analytical and numerical effects for a given pair

To illustrate how the long-term (analytical) and short-term (numerical) effects act on a given pair with different third-party species, Figure 1D shows the distribution of analytical and numerical system-level effects calculated on a hypothetical pair of symmetrically competing species *Ƶ* = {1, 2} (*a*_12_ = *a*_21_ = 0.22) when assembled together with different third species. Here, the interactions associated with the third species are chosen randomly from a normal distribution with *µ* = 0 and *σ* = 1 (all diagonal values are set to *a*_*ii*_ = 1). Note that for such multispecies systems there is generally a stable global equilibrium, and ℱ (𝒮) *>* 0. First, in this example, the projection contribution (first pink distribution) displays a large range of values including its maximum 1, confirming that the necessary conditions in (*θ*_1_, *θ*_2_) for pairwise coexistence in isolation are substantially expanded with the thirdparty species. Second, the long-term effects (second blue distribution) as well as the short-term effects (third orange distribution) show that the probability of pairwise coexistence with third-party species can either decrease or increase compared to the probability in isolation. Yet, both distributions are approximately centered at 1, especially for the analytical case (95% confidence intervals for the mean of analytical effects [0.956, 1.027]). In fact, Figure 1E shows that both effects are positively associated, but not perfectly correlated, as expected due to transient behavior. We have considered a wide variety of such systems and find that this property is similar for any pair of species under various numbers of randomly-assembled third-party species (Figs. S3-S4). This reveals that the probability of pairwise coexistence in isolation is very close to the expected probability across all possible (abiotic and biotic) conditions under both short-term and long-term behaviors. This result is confirmed and further illustrated by a perturbative analysis for weakly interacting systems in Sec. S5, where the center of the probability distribution of the analytical long-term effect varies from 1 only at fourth order in the interactions.

Lastly, for this particular pair of competing species, the buffering effects (fourth green distribution in Fig. 1D) are predominantly greater than 1, showing that the short-term effects tend to be stronger (relative to the long-term effects) with third-party species than in isolation. More generally, Figure 1F shows that negative (resp. positive) long-term effects of third-party species on pairwise coexistence are associated with a stronger (resp. weaker) buffering effect with third-party species than in isolation. In other words, short-term effects tend to be higher (resp. lower) than long-term effects if these long-term effects are less (resp. greater) than one. These results also hold for cases where the environmental effects are constrained to specific regions of the parameter space (Sec. S3, Fig. S6). All these results are also confirmed by the perturbative analysis for weakly interacting systems in Sec. S5.

### Comparing analytical and numerical effects across pairs

Next, we study how the long-term (analytical) and short-term (numerical) effects act across pairs with a given set of third-party species. Specifically, we randomly generate a five-dimensional interaction matrix **A**, where the interspecific effects (off-diagonal values) are sampled using a normal distribution with *µ* = 0 and *σ* = 0.25 (all diagonal values are set to *a*_*ii*_ = 1), making sure that the interaction matrix is diagonally dominant (and therefore globally stable). These values and conditions are chosen for illustrative purposes in line with experimental data in Sec. S1, but results are robust to this choice (Figs. 2 and S9). Additionally, to study how patterns can change as a function of the level of competition with third-party species, we generate a different five-dimensional interaction matrix, where the elements are the same as the first matrix except that all the off-diagonal signs are inverted. We measure the level of competition by the mean interaction strength. (We have also analyzed the association between the fraction of negative interactions and coexistence probability, but we did not find any systematic relationship; the reason is that even few strong negative interactions can generate relative negative impacts on pairwise coexistence.) Then, for each pair within the randomly-generated matrices **A**, we calculate the projection contribution *PC*(*Ƶ*, 𝒮), long-term effects *LE*(*Ƶ*, 𝒮), short-term effects *SE*(*Ƶ*, 𝒮), and the buffering effects *BE*(*Ƶ*, 𝒮). In order to calculate the simulated frequency of coexistence of each pair in isolation Sim(*Ƶ*), we run simulations at the pair level using the corresponding two-dimensional sub-matrices **A**[*i, j*; *i, j*] and sub-vectors ***θ***[*i, j*].

**Figure 2:**
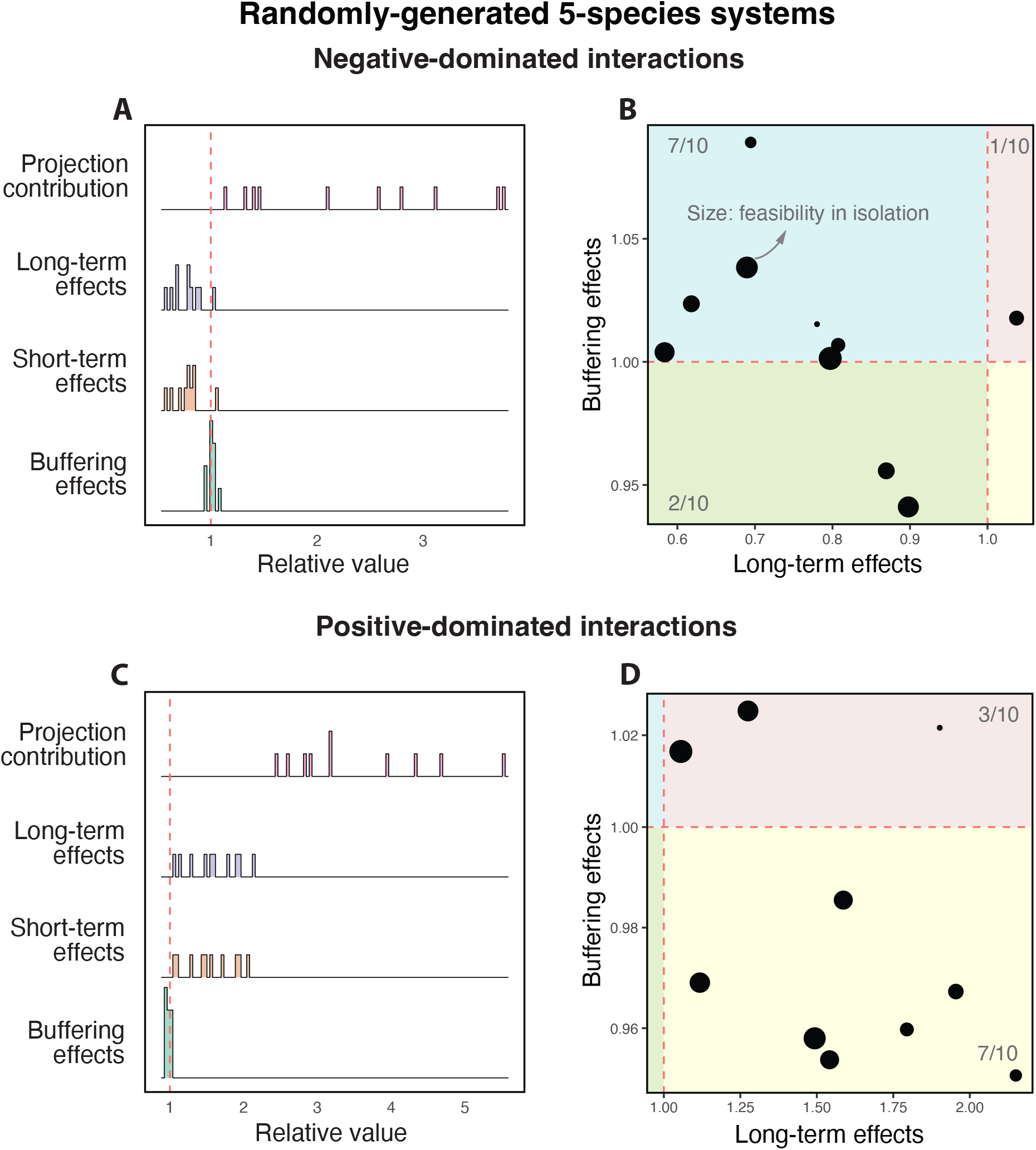
Cartographic representation of long-term and short-term differences among pairs with the same set of third-party species. Panels **A** and **C** show the system-level effects on 10 pairs within a randomly-generated five-species system characterized by negative-dominated and positive-dominated interactions, respectively (see text). Rows correspond to the projection distribution (*PC*(*Ƶ*, 𝒮)), long-term effects (*LE*(*Ƶ*, 𝒮)), short-term effects (*SE*(*Ƶ*, 𝒮)), and buffering effects (*BE*(*Ƶ*, 𝒮)), respectively. Note that each point in the distributions is a different pair within a set of third-party species. For reference, the dashed line shows the value of 1. Recall that the x-axis corresponds to the change in probability of pairwise coexistence with third-party species. Panels **B** and **D** illustrate the cartographic representation for all corresponding pairs based on beneficial (*LE*(*Ƶ*, 𝒮) > 1) or detrimental (*LE*(*Ƶ*, 𝒮) < 1) long-term effects and beneficial (*BE*(*Ƶ*, 𝒮) > 1) or detrimental (*BE*(*Ƶ*, 𝒮) < 1) short-term buffering effects. The size of points corresponds to the feasibility of pairs in isolation. The number of points in each region is annotated in gray. See Fig. S9 for a ten-species system.

## Results

Figures 2A-2B and 2C-2D summarize our results for a pair of negative-dominated (−⟨*a*_*ij*_⟩ = −0.051) and positive-dominated (−⟨*a*_*ij*_⟩ = 0.051) interaction matrices, respectively. Ecologically, negative-dominated interaction matrices correspond to primarily competitive systems. Instead, positive-dominated interaction matrices correspond to primarily cooperative systems. First, we found that the projection contribution (first pink distribution) is highly heterogeneous across pairs. Recall that this is a function of the combination between the pairwise interactions in *Ƶ* and the remaining interactions in the system 𝒮, where this combination changes from pair to pair. Second, we found that the long-term effects (second blue distribution) and short-term effects (third orange distribution) of third-party species can be disadvantageous (< 1) or beneficial (*>* 1) for pairs. Importantly, these effects are not equal across all pairs within the systems. Note that these distributions are only expected to be centered close to 1 for large enough systems given that each point in these distributions corresponds to a different pair with a fixed set of third-party species (not a fixed pair with different third-party species as in Fig. 1D). Third, the buffering effects (fourth green distribution) show values predominantly greater than (resp. less than) one in Figure 2A (resp. Figure 2C), revealing that the short-term effects tend to be stronger (resp. weaker) with third-party species than in isolation under negative-dominated (resp. positive-dominated) interactions. These strong (resp. weak) buffering effects are generated by the third-party species that also cause negative (resp. positive) effects over the long term.

To differentiate how long-term and short-term behaviors operate on each pair within a given set of third-party species, we construct a cartographic representation (38) of the relationship between long-term and buffering effects. Figures 2B and 2D show that the majority of pairs tend to be negatively (*LE* < 1) and positively (*LE >* 1) affected over the long-term within negative-dominated and positive-dominated random matrices, respectively. As a mirror image, the majority of pairs tend to be positively (*BE >* 1) and negatively (*BE* < 1) affected over the short-term within negative-dominated and positive-dominated random matrices, respectively. That is, on average, negative (resp. positive) long-term effects of third-party species on pairwise coexistence are typically linked with a stronger (weaker) short-term buffering effects with third-party species than in isolation, in agreement with the general results found in the previous section for a fixed pair in a variety of environments. Note that this cartographic representation can change as a function of the time span of the simulation (see Fig. S10). These effects were not correlated with pairwise coexistence in isolation (the size of points is not associated with the value of either axis).

### Experimental data

To test our theory in a more realistic setting, we use a publicly available data set of an *in vivo* gut microbial system. The data set (hereafter the fruit fly data set) was generated in Ref. (15) and it comprises a set of ten-day experimental trials of five interacting microbes commonly found in the fruit fly *Drosophila melanogaster* gut microbiota: *Lactobacillus plantarum* (Lp), *Lactobacillus brevis* (Lb), *Acetobacter pasteurianus* (Ap), *Acetobacter tropicalis* (At), and *Acetobacter orientalis* (Ao). This in vivo experimental study performed monoculture experiments for each species, co-cultures experiments for each pair, and poly-culture experiments for the quintet (Fig. 3 provides an illustration), as well as triplet and quartet experiments not relevant to our analysis here. Each experiment was replicated 48 times across different hosts and each species’ density was measured at the end of the observation period, providing one single record of persistence per replication for each species in monocultures, co-cultures, and poly-cultures (see Sec. S1 for details).

**Figure 3:**
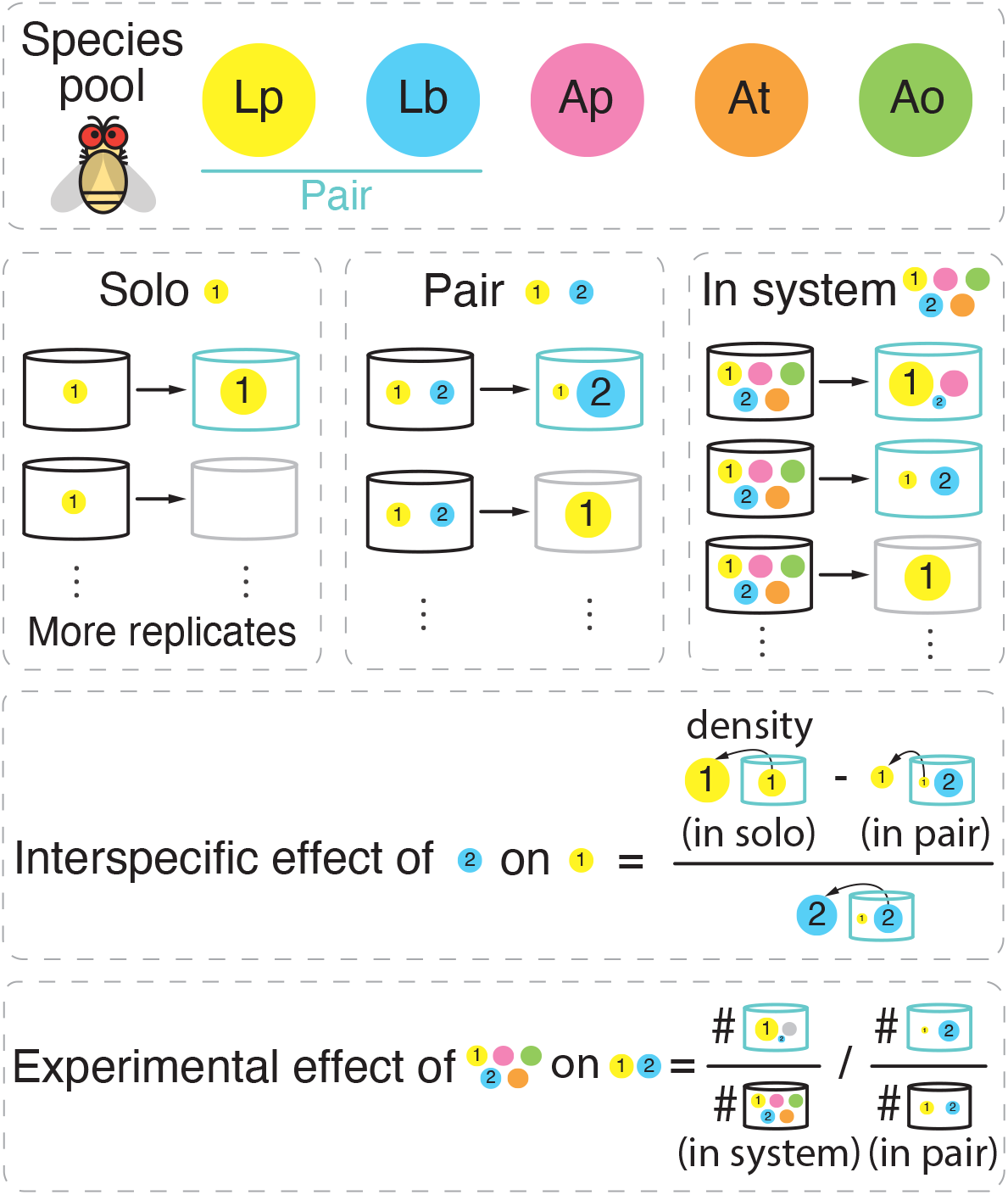
Illustration of experimental data, inference of pairwise interactions, and calculation of experimental effects. We use publicly available data of an *in vivo* gut microbial system. The data set was generated in Ref. (15) and it comprises ten-day experimental trials of five interacting microbes commonly found in the fruit fly *Drosophila melanogaster* gut microbiota: *Lactobacillus plantarum* (Lp), *Lactobacillus brevis* (Lb), *Acetobacter pasteurianus* (Ap), *Acetobacter tropicalis* (At), and *Acetobacter orientalis* (Ao). From the available experimental data, we used the results from monoculture experiments for each species, co-cultures experiments for each pair, and poly-culture experiments for the quintet. Each experiment was replicated across different hosts and each species’ density was measured at the end of the observation period, providing a single record of persistence for each species in monocultures, co-cultures, and poly-cultures per replication. The experimentally-inferred pairwise matrix **A** was inferred using the data of monoculture experiments for each species (e.g., species 1) and co-cultures experiments for pairs (e.g., pair {1, 2}) by evaluating the modified abundance of species 1 with and without species 2 at the end of the observation periods (see Sec. S3). We define the experimental effects of the third-party species on pair {1, 2} as the difference in the experimental frequencies of occurrence of the pair across all replicates with third-party species and the experimental frequencies of occurrence of the pair in isolation.

### Experimental system-level effects

To investigate the effects of third-party species (system-level effects) on pairwise coexistence using the *in vivo* data, for each pair *Ƶ*, we compute the observational frequency of coexistence with third-party species 𝒮 (i.e., within quintets) and in isolation *Ƶ* (i.e., within pairs) separately. To determine species coexistence, we classify a species statistically extinct in a trial if its relative abundance was less than 1% at the end of the observation period. We also test different extinction thresholds and obtain qualitatively the same results (Fig. S12). We define the *experimental effect* of third-party species on pairwise coexistence as the ratio between the frequency of coexistence with third-party species and the frequency in isolation (Fig. 3 provides an illustration):

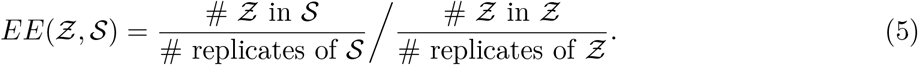

To compare the experimental and long-term (analytical) effects, we use the experimentally-inferred five-dimensional pairwise matrix **A** (see Sec. S2 for more details). This matrix was characterized by negative-dominant interactions (−⟨*a*_*ij*_⟩ = −0.078). Thus this experimental system is primarily competitive. Using this experimentally-inferred matrix, we analytically calculate the projected contributions *PC*(*Ƶ*, 𝒮) and long-term effects *LE*(*Ƶ*, 𝒮) of third-party species with |𝒮| = 5. We chose the quintet experiment (instead of the triplets or quartets) in order to have more pairs (10 with fives species) to perform statistics. In line with our numerical analysis, the *experimental buffering effect EBE*(*Ƶ*, 𝒮) is calculated by the ratio between the experimental effect *EE*(*Ƶ*, 𝒮) and long-term effect *LE*(*Ƶ*, 𝒮). We also construct a cartographic representation of the relationship between long-term effects and the experimental buffering effect in order to differentiate how long-term (analytical) and short-term (experimental) behaviors operate on each pair within the experimental system.

### Comparing analytical and experimental effects across pairs

We found that the experimental effects of third-party species on pairwise coexistence were very similar to those generated by simulations (Figs. 2, 4 and S11). Specifically, Figure 4A shows that the experimental multispecies system generated heterogeneous effects across pairs as illustrated by a wide distribution of projection contributions (first pink distributions), long-term effects (second blue distributions), and experimental effects (third orange distributions). Similar to simulations for systems with negative-dominant interactions, the *in vivo* experimental system shows predominately positive values of buffering effects (fourth green distributions). This shows that despite the negative long-term effects of the third-party species on pairwise coexistence, there is a stronger buffering effect with third-party species than in isolation. Focusing on the cartographic representation of differences among pairs, Figure 4B shows that the majority of pairs tend to be negatively affected over the long-term (*LE* < 1) and positively affected over the short-term (*EBE >* 1)—consistent with the results obtained for negative-dominant matrices (Fig. 2B). This cartographic representation can also help to understand specific pairwise cases. For example, the pair (Ap, At) was observed with negative long-term effects but with a strongly positive buffering effect over the short-term (top left region). In contrast, the pair (Lp, Ap) exhibited the complete opposite effects. In turn, the pair (Ap, Ao) exhibited positive effects over both the long-term and short-term, suggesting that this pair is substantially benefited by the third-party species. This classification can change as a function of the experimental details. Importantly, these effects were again uncorrelated with pairwise coexistence in isolation (the size of points in Fig. 4B is not associated with the value of either axis), confirming the importance of third-party-species dynamics on pairwise coexistence.

**Figure 4:**
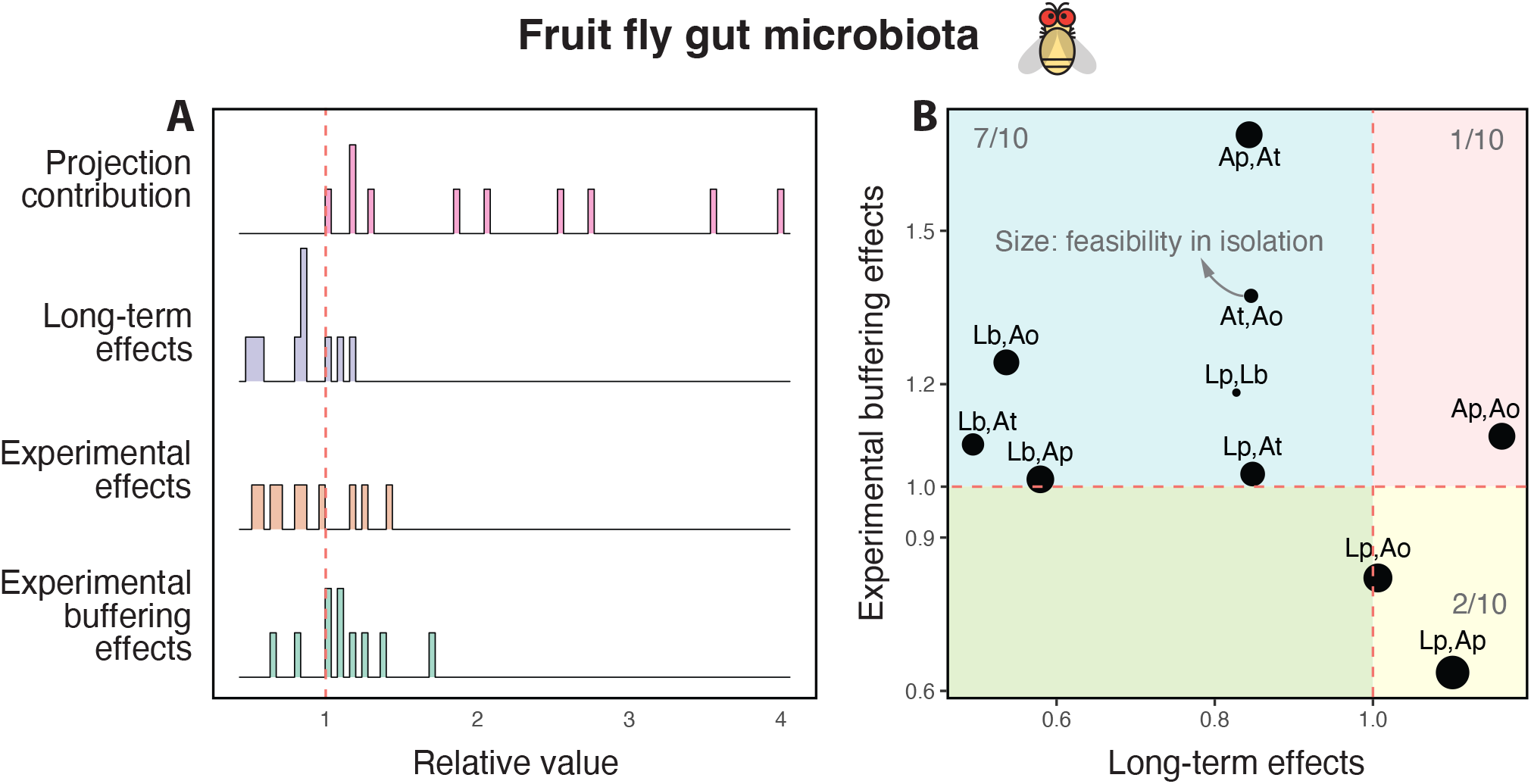
Comparing system-level effects among pairs within a five-species *in vivo* experimental system. This figure shows similar information to Fig. 2, but using the experimental data and the experimentally-inferred interaction matrix. For each of the 10 pairs in the experimental fruit fly system, Panel **A** shows the projection distribution *PC*(*Ƶ*, 𝒮) (first row pink distribution), long-term effects *LE*(*Ƶ*, 𝒮) (second row blue distribution), experimental effects *EE*(*Ƶ*, 𝒮) (third row orange distribution), and experimental buffering effects *EBE*(*Ƶ*, 𝒮) (fourth row green distribution). Panel **B** shows the cartographic representation of how third-party species affect the long-term and short-term behavior of each of its constituent pairs. Long-term effects can be beneficial (*LE*(*Ƶ*, 𝒮) *>* 1) or detrimental (*LE*(*Ƶ*, 𝒮) < 1). In this experimental system, the majority of short-term effects tend to be beneficial (*EBE*(*Ƶ*, 𝒮) < 1) in line with negative-dominated interaction matrices (Fig. 2B). See text or Fig. 3 for all species names.

## Discussion

A major challenge in ecological research has been understanding how species interactions and diverse environments affect the opportunities for species coexistence (2, 4). Many research efforts have been devoted to studying the coexistence of either two species in isolation or that of a particular set of multiple species (8). Yet, species are seldom in isolation and the coexistence of a specific combination of multiple species is expected to be rare in a random environment (5, 24, 25). This has prompted us to better understand the extent to which the coexistence of a particular pair or presumed community of species can change between being in isolation to being part of a larger multispecies system. The main problem resides in the fact that in natural settings, most of the abiotic and biotic effects acting on species are unknown. As a response to this problem, we have introduced an analytical formula to estimate the extent to which third-party species can affect the probability that a particular pair of species coexist under environmental uncertainty (all possible conditions are equally likely to happen). This formula is based on information about pairwise interactions from a given pool of species and assumes that the total effect of all additional biotic and abiotic factors can be captured by a phenomenological random parameter for each species (i.e., the effective growth rate parameter *θ*). This kind of question is also a critical focus and challenge in bio-restoration and bio-medicine, where it is unclear whether a large system can have different effects on its different pairs of species (6, 7, 48–50). In restoration, for example, it may be that in a given ecological context where the species composition has been drastically modified or reduced, a given pair of species that could coexist in the original full ecological community may no longer have a feasible equilibrium in the modified species composition. In such a situation, better understanding the kind of analytic effects described here could help inform restoration ecologists regarding which other species must simultaneously be reincorporated to enable a specific pair to flourish.

Following our analytical framework, we have shown that the diversity of multispecies systems opens the opportunity for pairwise coexistence regardless of whether a given pair can coexist or not in isolation. This phenomenon occurs since the environmental conditions limiting the coexistence of a pair in isolation (parameter values acting on a plane) are replaced by a new set of conditions acting on all species, whose projection onto the two-dimensional space tends to cover the pair’s entire plane. Thus, the possibility of pairwise coexistence with third-party species only depends on the new set of conditions in the multidimensional space of the entire system. The magnitude of projection reflects the range of compatible environments for the pair to coexist. This can be interpreted as the extent to which third-party species can act as ecosystem engineers (26, 27) and modify the environment into more suitable habitats. Yet, the probability of pairwise coexistence within larger systems still depends on the effective growth rates associated with third-party species. The long-term effects measure the difference in the probability of pairwise coexistence with and without third-party species. This can be interpreted as the relative benefit (or harm) of third-party species on pairwise coexistence. Importantly, under environmental uncertainty, the expected value of the distribution of probabilities generated by all possible multispecies systems is approximately the same as the probability in isolation. That is, we can usefully approximate the likelihood of two species coexisting in diverse assemblages (or diverse environmental conditions) based on the understanding of their pairwise interactions. We have shown analytically that the deviation of the mean of the distribution from the isolated probability scales as *a*^4^ for simple model systems with interactions of order *a*, and confirmed numerically that this effect is still very small for larger systems outside the analytic framework. This reasoning can be applied to any subset of species embedded in large multispecies systems.

While analytical measures are useful for increasing our general understanding of pairwise coexistence, the assumptions behind these measures may never be met in real-world systems. For example, our analytical measures are based on the long-term asymptotic equilibria achieved by systems in arbitrary environmental conditions. Yet, both simulations and experiments are always restricted to or biased by short-term (finite) effects. Hence, to provide an additional understanding of such short-term effects, we have compared our analytical expectations against numerical and experimental outcomes. As expected, we confirmed that the probability of pairwise coexistence over the short term (numerically and experimentally) is greater than pairwise coexistence over the long term (12). Yet, it has been unclear whether this buffering effect should be equal with third-party species and in isolation. We have found that this increase is typically greater (resp. smaller) for pairs with third-party species than in isolation if the third-party species affect the pairs negatively (resp. positively) in the long term. In other words, we have found that when third-party species tend to reduce (resp. increase) the coexistence probability of a pair, they tend to exhibit slower (resp. faster) rates of competitive exclusion. This illustrates potential differences between what we may see in experiments and the expected long-term dynamics of ecological systems. Moreover, these effects are not equal across all pairs within a system. Importantly, we have shown that we can study these differences using a cartographic representation of how third-party species are expected to affect the long-term and short-term behavior of each pair.

As a final note, it is worth mentioning that our theory assumes that environmental conditions are uniformly distributed across the entire parameter space. Instead, under experimental settings, much stronger constraints on environmental conditions can be expected. This implies that the probability of pairwise coexistence should be different between theoretical and experimental analyses. Yet, these experimental constraints should also be expected to operate on pairs with third-party species and in isolation. Therefore, system-level effects (ratio between the probability in multispecies systems and in isolation) should be comparable between theoretical and experimental analyses. Indeed, our theoretical analysis (Fig. 2) showed similar patterns to the experimental analysis (Fig. 4), suggesting that our phenomenological framework is applicable to real-world settings and can be used to increase our understanding of long-term and short-term ecological dynamics.

## Acknowledgments

We thank the Editor Sara Mitri, Aming Li, and two anonymous reviewers for their highly constructive comments that led to the improvement of this work. S.S. was supported by NSF DEB-2024349. W.T. was supported in part by a grant from the Schmidt Futures Foundation.

## Notes

**Data accessibility** The code supporting the results is novel and will be archived on Zenodo upon publication. This code is available to reviewers on Github: https://github.com/MITEcology/Deng_2022_1

### Competing Interest Statement

The authors have declared no competing interest.

### Summary of Updates

Title revised; minor revisions in the text.

