## Supplemental Materials for "Understanding the impact of third-party species on pairwise coexistence"

#### Contents

**S1 Experimental communities: Fruit fly gut microbiota** **S2**

**S2 Inferring the interaction matrix** **S3**

**S3 An example of constrained environmental conditions** **S4**

**S4 Monte Carlo estimate of measures $F(\mathcal{C}, \mathcal{S})$** **S7**

**S5 A simple analytic model** **S7**

**S6 Supplementary Figures** **S12**

### S1 Experimental communities: Fruit fly gut microbiota

The studied experimental community is compiled from Ref. (15). For the readers' convenience, below we summarize information about the species, medium, experiments, and measurements as written in Ref. (15), the authors of which performed the experiments. Please refer to Ref. (15) for more details.

- Host species: Wolbachia-free and virus-free Drosophila melanogaster Canton-S flies.
- Medium: 6.67% cornmeal, 2.7% active dry yeast, 1.6% sucrose, 0.75% sodium tartrate, 0.73% ethanol, 0.68% agar, 0.46% propionic acid, 0.09% methylparaben, 0.06% calcium chloride, and 0.01% molasses.
- Experimental conditions: 25 °C, 60% humidity, 12:12 h light:dark cycles, sterile conditions.
- Bacterial strains: Five fermentative lactic acid bacteria and acetic acid bacteria commonly occurring in the wild fly gut. Specifically, the five bacteria are
  - *Lactobacillus plantarum* (Lp),
  - *Lactobacillus brevis* (Lb),
  - *Acetobacter pasteurianus* (Ap),
  - *Acetobacter tropicalis* (At),
  - *Acetobacter orientalis* (Ao).
- Bacterial abundance calculations: On the $10^{\text{th}}$ day of inoculation, flies were washed in 70% ethanol before being bead-beaten in 96-well plates with a custom-made attachment. Lysates were pinned onto selective media using a 96-pin replicator (Boekel), visually scored, and colony-forming units (CFUs) were enumerated.
- Bacterial combinations: 5 single bacterium, 10 pairs of bacteria, 10 triplets of bacteria, 5 quartets of bacteria, 1 quintet of all bacteria.
- Replicates: 48 replicates were performed for each bacterial combination.
- Gnotobiotic preparation: Per fly vial, each bacterium was prepared $5 \times 10^6$ CFUs (50 $\mu L$ of $10^8$ bacteria per milliliter) and then mixed according to the bacterial combinations. Germ-free flies were sorted into these vials.

The results of these experiments were datasets consisting of the abundances of the different bacteria species in each of the combinations and replicates after 10 days.

#### S2 Inferring the interaction matrix

To infer the pairwise interaction matrix from the fruit-fly experiments (15), we considered a subset of the above data sets containing only one or two species. Therefore, the available information indicates the abundance of species $i$ in isolation, and how its abundance is modified when species $j$ is included. A small (resp. large) change in the density of species $i$ due to the inclusion of species $j$ is evidence of a negligible (resp. significant) $a_{ij}$. To quantify the value of $a_{ij}$, let $x_i^*$ denote the equilibrium density of the $i$-th species in isolation. Let $(y_i^*, y_j^*)$ denote the equilibrium density of species $i$ and $j$ when both are present in the experiment (co-cultures). Then, for all data such that $x_i^* \neq 0$ and $y_i^* \neq 0$, the LV model in this form:

$$\frac{\mathrm{d}N_i}{\mathrm{d}t} = N_i \left( \theta_i - \sum_{j=1}^{S} a_{ij} N_j \right) \tag{S1}$$

implies that the following equations must be satisfied:

$$\begin{aligned} \theta_i - a_{ii} x_i^* &= 0, \\ \theta_i - a_{ii} y_i^* - a_{ij} y_j^* &= 0. \end{aligned} \tag{S2}$$

Note that, since only steady-state measurements are available, there are fundamental limitations in the information of $a_{ij}$ that can be inferred (32). Namely, in the above equations for each species $i$ we have three unknown $(\theta_i, a_{ii}, a_{ij})$ but only two equations, implying there is not sufficient information to constrain the three unknowns. To circumvent this limitation, we can use the fact that the variables are homogeneous measures and any normalization will not affect the outcomes, such as assuming $\hat{a}_{ii} = 1$ (51). With this assumption, and subtracting the first from the last line of Eq. (S2), we obtain

$$a_{ij} = \frac{x_i^* - y_i^*}{y_j^*}, \tag{S3}$$

which can be used to estimate $a_{ij}$ from the experimental data.

Importantly, since there are several replicates for $x_i^*$ and $(y_i^*, y_j^*)$, Eq. (S3) actually characterizes a distribution of possible values for $a_{ij}$ obtained by taking an arbitrary pair of replicates. To obtain the single value necessary for our predictions, we defined

$$\hat{a}_{ij} = \mathrm{Median}(a_{ij}), \quad i \neq j, \tag{S4}$$

estimated using a Bootstrap method with all possible $48 \times 48$ pairs. By repeating this process for all pairs of species, we obtained the following estimate for the interaction matrix:

$$\hat{A} = \begin{array}{c} \\ \text{Lp} \\ \text{Lb} \\ \text{Ap} \\ \text{At} \\ \text{Ao} \end{array} \begin{array}{c} \begin{array}{ccccc} \text{Lp} & \text{Lb} & \text{Ap} & \text{At} & \text{Ao} \end{array} \\ \begin{pmatrix} 1 & 0 & -0.182 & -0.091 & -0.197 \\ 1.119 & 1 & 0.097 & 0.34 & 0.094 \\ -0.592 & 0 & 1 & 0 & 0 \\ 0.288 & 0.036 & 0 & 1 & 0.576 \\ 0 & 0 & 0 & 0.239 & 1 \end{pmatrix} \end{array}. \tag{S5}$$

Fig. S11 shows the accuracy of this pairwise matrix in predicting the multispecies dynamics.

### S3 An example of constrained environmental conditions

In the main text, we assume the environmental conditions are random and heterogeneous so that the effective growth rates $\theta$ are uniformly distributed on the closed unit sphere (Fig. 1). Yet, real ecological systems and experiments may suggest a specific range of environmental variation (e.g., *in vitro* laboratory experiments are typically highly-controlled), which geometrically constrains the parameter space from the entire unit sphere to a specific region. To illustrate how our framework continues to operate under these constrained scenarios, here we calculated the system-level effects for the fixed pair $\{1, 2\}$ in Fig. 1D within random 3-dimensional systems under constrained environments. Specifically, we assume that the effective growth rate of individual species ranges between $[-0.9, 1]$ due to environmental constraints. Figure S1 illustrates our analytical system-level effects (Fig. S1A) and the projection contribution (Fig. S1B) in the constrained parameter space.

Note that the long-term (analytical) effects are evaluated in 3-dimensional space, and the projection contribution is in 2-dimensional space. Besides, all the calculations of constrained effects can be based on the effects without constraint. The feasibility $F(\mathcal{Z})$ of the fixed pair $\mathcal{Z} = \{1, 2\}$ in isolation without constraint is 0.18, and the corresponding interaction matrix is $\begin{pmatrix} 1 & 0.22 \\ 0.22 & 1 \end{pmatrix}$.

Specifically, for the long-term effects (Fig. S1A), we can firstly solve the coordinates of the two intersection points: M $(0.43, 0.093, -0.9)$, N $(0.093, 0.43, -0.9)$. The plane $\theta_3 = -0.9$ removes the region below it, whose hypervolume (i.e., surface area) is 0.093, from the feasibility regions $P(\mathcal{Z}, \mathcal{S})$ of the pair $\mathcal{Z} = \{1, 2\}$ within a 3-species system $\mathcal{S}$ (i.e., pink region and blue region). Besides, the three planes $\theta_1 = -0.9,\ \theta_2 = -0.9,\ \theta_3 = -0.9$ each eliminate equal hypervolumes, which is 0.52, from the entire unit sphere. Then we have the hypervolume of the constrained parameter space $4\pi - 3 \times 0.52 = 11.01$. Thus, the constrained feasibility $P'(\mathcal{Z}, \mathcal{S})$ of the pair $\mathcal{Z} = \{1, 2\}$ within a 3-species system $\mathcal{S}$ can be computed by

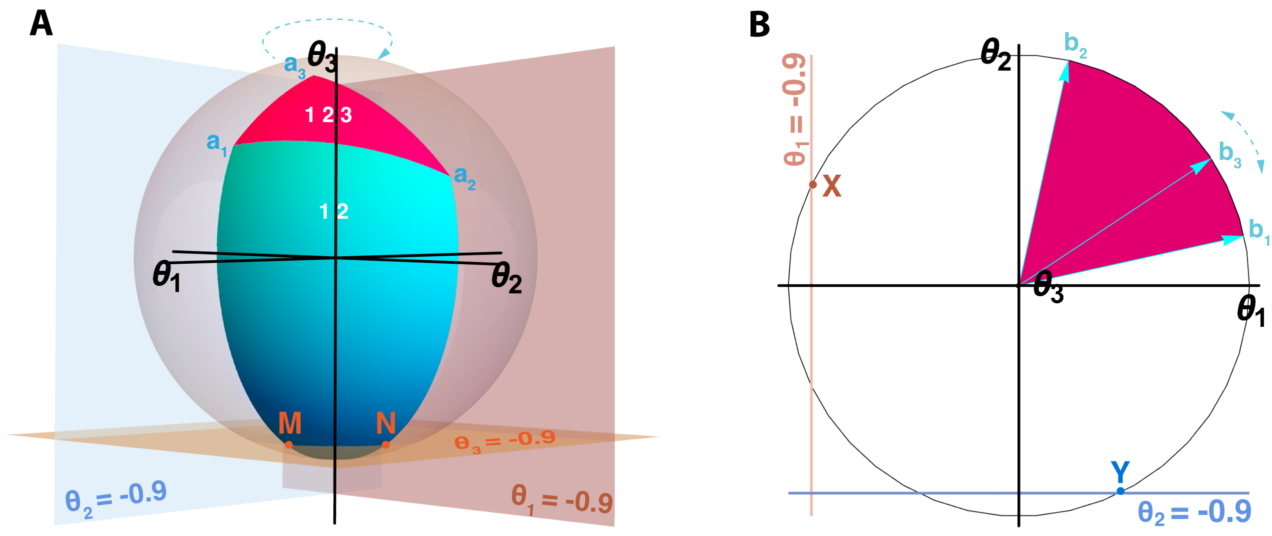


Supplementary Figure S1: **Illustration of long-term effects and projection contribution under constrained environments.** Panel **A** shows the constrained feasibility of the pair within a random 3-species system. The spanning vectors $\mathbf{a}_1, \mathbf{a}_2$ and $\mathbf{a}_3$ are the three column vectors of the interaction matrix. The parameter space is constrained by three planes $\theta_1 = -0.9, \theta_2 = -0.9$ and $\theta_3 = -0.9$. Points M and N are the intersection points of the colored feasibility region and the plane $\theta_3 = -0.9$. Panel **B** shows the constrained projection in 2-dimensional space. The spanning vectors $\mathbf{b}_1, \mathbf{b}_2$ and $\mathbf{b}_3$ are the projections of $\mathbf{a}_1, \mathbf{a}_2$ and $\mathbf{a}_3$, respectively. The parameter space is constrained by two lines $\theta_1 = -0.9$ and $\theta_2 = -0.9$. Points X and Y are two intersection points of the unit circle and the two lines.

$$P'(\mathcal{Z}, \mathcal{S}) = \frac{P(\mathcal{Z}, \mathcal{S}) \times 4\pi - 0.093}{11.01},$$

where $P(\mathcal{Z}, \mathcal{S})$ is the feasibility of the pair $\mathcal{Z} = \{1, 2\}$ within a 3-species system $\mathcal{S}$ without constraint.

Similarly, the constrained feasibility $F'(\mathcal{Z})$ of the pair $\mathcal{Z} = \{1, 2\}$ in isolation can be computed by

$$F'(\mathcal{Z}) = \frac{F(\mathcal{Z}) \times 4\pi - 0.093}{11.01} = \frac{0.18 \times 4\pi - 0.093}{11.01} = 0.20,$$

where $F(\mathcal{Z})$ is the feasibility of the pair $\mathcal{Z} = \{1, 2\}$ in isolation without constraint.

Based on the definition, we can obtain the long-term effects of any random 3-species system $\mathcal{S}$ on pair $\mathcal{Z} = \{1, 2\}$ under constrained environments by

$$LE(\mathcal{Z}, \mathcal{S}) = \frac{P'(\mathcal{Z}, \mathcal{S})}{F'(\mathcal{Z})}.$$

For the projection contribution (Fig. S1B), it is clear that the projection region of a 3-species system spanned by $\mathbf{b}_1$, $\mathbf{b}_2$ and $\mathbf{b}_3$ is always larger or equal to the feasibility region of the pair in isolation spanned by $\mathbf{b}_1$ and $\mathbf{b}_2$. The two lines $\theta_1 = -0.9$, $\theta_2 = -0.9$ each eliminate equal hypervolumes (arc length) from the unit circle. By the Pythagorean theorem, we can calculate the angle and then get the hypervolume 0.90 of single elimination. Thus, the constrained feasibility $F''(\mathcal{Z})$ of the pair $\mathcal{Z} = \{1, 2\}$ in isolation is

$$F''(\mathcal{Z}) = \frac{F(\mathcal{Z}) \times 2\pi}{2\pi - 2 \times 0.90} = \frac{0.18 \times 2\pi}{2\pi - 2 \times 0.90} = 0.25,$$

where $F(\mathcal{Z})$ is the feasibility of the pair $\mathcal{Z} = \{1, 2\}$ in isolation without constraint.

Note that the vector $\mathbf{b}_3$ corresponding to a random species can move freely along the unit circle. If the projection region is not the full two-dimensional parameter space, then its hypervolume is maximized when $\mathbf{b}_3$ moves to the point X $(-0.9, 0.44)$ or Y $(0.44, -0.9)$. At these two points, the constrained projection $\text{Proj}'(\mathcal{Z}, \mathcal{S})$ is

$$\text{Proj}'(\mathcal{Z}, \mathcal{S})_\text{X} = \text{Proj}'(\mathcal{Z}, \mathcal{S})_\text{Y} = \frac{\pi - 0.90/2 - (\frac{\pi}{2} - F(\mathcal{Z}) \times 2\pi)/2}{2\pi - 2 \times 0.90} = \frac{\pi - 0.45 - 0.22}{2\pi - 2 \times 0.90} = 0.55.$$

Otherwise, if the projection region covers the entire constrained 2-dimensional space, then the $\text{Proj}'(\mathcal{Z}, \mathcal{S}) = 1$.

In general, when $\mathbf{b}_3$ is moving on the (longer) arc between points X and Y, the constrained projection $\text{Proj}'(\mathcal{Z}, \mathcal{S})$ can be computed by

$$\text{Proj}'(\mathcal{Z}, \mathcal{S}) = \frac{\text{Proj}(\mathcal{Z}, \mathcal{S}) \times 2\pi}{2\pi - 2 \times 0.90} = \frac{\text{Proj}(\mathcal{Z}, \mathcal{S}) \times 2\pi}{4.48} \in [0.25, 0.55] \cup \{1\},$$

where $\text{Proj}(\mathcal{Z}, \mathcal{S})$ is the projection without constraint.

Hence, according to the definition, the amount of projection contribution increasing the feasibility region in the constrained parameter space is

$$PC(\mathcal{Z}, \mathcal{S}) = \frac{\text{Proj}'(\mathcal{Z}, \mathcal{S})}{F''(\mathcal{Z})}.$$

For the short-term (simulated) effects, we sample the points on the entire unit sphere, remove the ones located outside of the constrained region, and then calculate the frequency of pairwise coexistence based on the remaining points.

The buffering effect can be easily obtained by the ratio between the short-term and long-term effects.

Figure S6 shows the distributions of aforementioned system-level effects for the fixed pair $\{1, 2\}$ within 50 different 3-dimensional systems under constrained environmental conditions. This shows that short-term effects tend to be higher (resp. lower) than long-term effects if these long-term effects are less (resp. greater) than one—confirming that our main results also hold for cases where the environmental effects are constrained to specific regions of the parameter space.

#### S4 Monte Carlo estimate of measures $F(\mathcal{C},\mathcal{S})$

One way to efficiently estimate the measure $F(\mathcal{C},\mathcal{S})$ of a given community $\mathcal{C}$ within a larger multispecies system $\mathcal{S}$ is to use a simple Monte Carlo approach. We can randomly sample a point on the unit sphere in $S = |\mathcal{S}|$ dimensions by picking a random vector $\mathbf{x} = \sum_{i \in \mathcal{S}} \nu_i \mathbf{e}_i$ where $\nu_i$ is chosen from a Gaussian distribution with a common variance for each $i$. We can efficiently test whether $\mathbf{x} \in D_F(\mathcal{C},\mathcal{S})$ by checking $S$ linear conditions of the form $\mathbf{x} \cdot \mathbf{b} > 0$, as described below. We can then rapidly accumulate statistics on the fraction of random points in the sphere that lie in $D_F(\mathcal{C},\mathcal{S})$ to estimate $F(\mathcal{C},\mathcal{S})$.

In further detail, the linear conditions that determine which community $\mathcal{C} \subset \mathcal{S}$ is associated with a given point $\mathbf{x}$ can be described in terms of the boundaries between a community $\mathcal{C}$ and a community $\mathcal{C}' = \mathcal{C} \cup \{i\}, i \notin \mathcal{C}$. A necessary condition for $\mathbf{x} \in D_F(\mathcal{C},\mathcal{S})$ or $\mathbf{x} \in D_F(\mathcal{C}',\mathcal{S})$ is that

$$(\mathsf{det}[\mathbf{a}'_{j_1} \ldots \mathbf{a}'_{j_C} \mathbf{x}]) = \mp(\mathsf{det}[\mathbf{a}'_{j_1} \ldots \mathbf{a}'_{j_C} \mathbf{e}_i]),\ j_k \in \mathcal{C}, \tag{S6}$$

where $\mathbf{a}'$ denotes the restriction of the vector $\mathbf{a}$ to the subspace associated with community $\mathcal{C}'$, and the negative/positive sign is associated with the community $\mathcal{C}/\mathcal{C}'$. This follows because the linear hypersurface that separates $\mathcal{C}, \mathcal{C}'$ is spanned by the vectors $\mathbf{a}'_{j_k}$ and $\mathbf{e_i}, i \notin \mathcal{C}'$. (Note that these conditions continue to be valid in the degenerate case $\mathcal{C} = \{\}$, which is needed to fix the boundaries of communities $\mathcal{C}' = \{i\}$ containing only a single species.) To check whether $\mathbf{x} \in D_F(\mathcal{C},\mathcal{S})$, we thus simply check each of the $S$ conditions associated with the boundaries of $\mathcal{C}$ with communities that differ by a single species $i$, where $i$ is added or subtracted from the community depending on whether $i \in \mathcal{C}$. If all these conditions are satisfied then $\mathbf{x} \in D_F(\mathcal{C},\mathcal{S})$.

Since each of the determinants on the LHS of (S6) can be written in the form

$$\mathsf{det}[\mathbf{a}'_{j_1} \ldots \mathbf{a}'_{j_C} \mathbf{x}] = \mathbf{b}_{C,i} \cdot \mathbf{x}\,,$$

by computing the vectors $\mathbf{b}_{C,i}$ in advance for each combination $C, i$ we can efficiently sample and test many points to give a good Monte Carlo estimate of $F(\mathcal{C},\mathcal{S})$. A simple mathematica code implementing this Monte Carlo algorithm is available with the other code supporting the results of this work.

#### S5 A simple analytic model

*Simple analytic model:*

In this section we briefly describe a simple model that illustrates some of the methods and results of the main paper in a simplified situation amenable to direct analytic treatment. We consider a fixed system ($\mathcal{Z}$) of two species in a larger system ($\mathcal{S}$) containing a third species where the total interaction matrix is parameterized by

$$\mathbf{A} = \begin{pmatrix} 1 & \tan\phi_1 & a \\ \tan\phi_2 & 1 & b \\ c & d & 1 \end{pmatrix} . \tag{S7}$$

In some of the following analysis we take all interactions with the third species to be small, on the order of a small parameter $\epsilon$, so $a = \tilde{a}\epsilon, b = \tilde{b}\epsilon, etc.$, and perform a perturbative expansion in $\epsilon$. We also set $\phi_1 = \phi_2 = \phi$ for simplicity in some of the analysis, corresponding to a symmetric interaction $a_{12} = a_{21}$.

While the focus here is on systems of three species, some of the perturbative results are directly relevant for systems with more species.

*Projection contribution:*

We consider the computation of the projection contribution (1), governing the range of conditions $(\theta_1, \theta_2)$ under which it is *possible* that species 1 and 2 can coexist (i.e., coexistence is possible for some $\theta_3$). The angle subtended by the vectors $(1, \tan\phi), (\tan\phi, 1)$ in the 2D system is $\alpha = \pi/2 - 2\phi$, so

$$F(\mathcal{Z}) = \frac{\alpha}{2\pi} = 1/4 - \phi/\pi \, .$$

The projection region $D_{\text{proj}}(\mathcal{Z}, \mathcal{S})$ only depends upon $a, b$. Parameterizing $a = \epsilon\cos\eta, b = \epsilon\sin\eta$, from simple geometry we can compute the projection fraction

$$\text{Proj}(\mathcal{Z}, \mathcal{S}) = \begin{cases} 1/4 - (\eta + \pi)/2, & -\pi/2 - \phi \leq \eta \leq \phi \\ 1/4 - \phi/\pi, & \phi \leq \eta \leq \pi/2 - \phi \\ (\eta - \phi)/2\pi, & \pi/2 - \phi \leq \eta \leq \pi + \phi \\ 1, & \pi + \phi \leq \eta \leq 3\pi/2 - \phi \end{cases}$$

Thus, we see that for random $a, b$ the distribution of the projection fraction is such that Proj $=$ $1/4 - \phi/\pi$ with probability $1/4 - \phi/\pi$, Proj $= 1$ with probability $1/4 - \phi/\pi$, and uniformly distributed between the minimum value and $1/2$ in the remaining cases of total probability $1/2 + 2\phi/\pi$. The projection contribution $PC$ arises after dividing this by the minimum projection value $1/4 - \phi/\pi$. This matches qualitatively with, e.g., Figure 1 D, where there are similar contributions to PC at 1 and $1/F(\mathcal{Z})$, and a roughly uniform distribution between 1 and $1/2F(\mathcal{Z})$. Note that this analysis is valid for arbitrary $\epsilon$, not necessarily small.

*Long-term system effects*

We now consider the long-term (analytic) effect governing the *probability* that the species 1, 2 can coexist for randomly chosen environmental conditions $\theta_1, \theta_2, \theta_3$. This probability is given by the union of the feasibility regions $P(\mathcal{Z}, \mathcal{S}) = F(\{1, 2\}, \mathcal{S}) + F(\{1, 2, 3\}, \mathcal{S})$. We can analyze this analytically using an analytic formula for the solid angle $\Omega$ of a triangle bounded by three rays $\mathbf{x}, \mathbf{y}, \mathbf{z}$

$$\tan(\Omega) = \mathsf{det}(\mathbf{xyz})/(xyz + (\mathbf{x}\cdot\mathbf{y})z + (\mathbf{x}\cdot\mathbf{z})y + (\mathbf{y}\cdot\mathbf{z})x)\,.$$

To leading order in $\epsilon$, for fixed $a, b, c, d$ of order 1, we have

$$P(\mathcal{Z}, \mathcal{S}) = \frac{1}{4\pi}\left(2\alpha - (a+b)(\cos\phi - \sin\phi) + \mathcal{O}(\epsilon^2)\right) = F(\mathcal{Z})(1 - (a+b)(1 - \alpha^2/24 + \mathcal{O}(\alpha^4))/2\sqrt{2})\,,$$

so to this order the long-term effect is $\mathrm{LE}(\mathcal{Z}, \mathcal{S}) = 1 - (a+b)/2\sqrt{2} + \mathcal{O}(\epsilon^2, \alpha^2)$ for small angles $\alpha$, as can be seen naturally from the geometry. On the other hand, separating the parameters $\phi_1, \phi_2$ and taking both to be small of order $\epsilon$, we have $\alpha \sim \pi/2 - \phi_1 - \phi_2$, so $\mathrm{F}(\mathcal{Z}) \sim 1/4 - (\phi_1 + \phi_2)/\pi$, and

$$\mathrm{LE}(\mathcal{Z}, \mathcal{S}) = 1 - (a+b)/\pi + \mathcal{O}(\epsilon^2) \sim 1 - 0.318(a+b)$$

This shows that to leading order for a symmetric probability distribution on $a + b$ around 0, the distribution on LE is symmetric and centered around 1, compatible with, e.g., Figure 1 D. A more careful analysis gives the second order effect

$$F(\mathcal{Z}, \mathcal{S}) = 2\alpha - (a+b)(\cos\phi - \sin\phi) + \frac{1}{2}\left(bc + ad - (ac + bd)\cos\alpha + (2ab + bc + ad)\sin\alpha\right) + \mathcal{O}(\epsilon^3)\,. \tag{S8}$$

Since there are no quadratic terms in any of the coefficients $a, b, c, d$, in a random distribution where these coefficients are uncorrelated there is also no second order shift in the mean of the LE distribution. All cubic terms are odd in at least one variable so also do not shift the mean in a symmetric distribution. There are, however, quartic terms, such as $-a^4 \cos\alpha \sin\alpha/8$ that contribute to a shift of the mean. This gives some indication, however, of the reason why the mean of LE is so close to 0, so that the presence of a random environment has on average negligible effect on the probability of coexistence of a fixed pair.

Note that the linear effect of small interactions of a third species on LE for a given pair of species carries over to larger numbers of additional species; the linear effect on LE of a set of $|\mathcal{S}| - 2$ additional species on a given pair is simply given by the sum of the linear effects from each individual additional species. Thus, to leading order, $\mathrm{LE}(\mathcal{Z}, \mathcal{S}) = \prod_{i=3}^{|\mathcal{S}|} \mathrm{LE}(\mathcal{Z}, \mathcal{Z} \cup \{i\})$.

*Short-term effects*

A detailed analysis of the short-term effects associated with changes in the time of persistence of the pair in the larger community is somewhat more subtle, even in this simple analytic model, as it involves dynamics. To get a simple sense of the order of magnitude of this effect, however, we can consider the dynamical equations such as

$$\dot{N}_1 = N_1(\theta_1 - N_1 - \tan\phi_1 N_2 - aN_3) \tag{S9}$$

from the interaction matrix (S7), where in the analysis of this section we take $\phi_1, \phi_2$ small so $\tan\phi_i \sim \phi_i$. We set all the parameters in (S7) to vanish except $a$ and focus on the effect of a small interaction term of this type; the parameters other than $a$ and the equivalent parameter $b$ do not have effects at linear order. For a fixed threshold of extinction, we can associate the extinction boundary in $\theta_1$ with a certain effective negative growth rate

$$\theta_1 - N_1 - aN_3 < \text{constant}\,. \tag{S10}$$

The change in this boundary value of $\theta_1$ under the perturbation $a$ will be of order $aN_3 \sim a\theta_3$. Thus, when the extinction threshold $\eta$ is fixed and very small and the time of simulation $T$ is long enough that the extinction boundary is near the coexistence boundary ($T \gg |\ln\eta|$), the region in the $\theta$-sphere where the extinction condition on the pair in isolation and in the larger set differ will be near the boundary $\theta_1 = 0$ and will have an area of roughly $a\int_0^{\pi/2}\sin\theta = a$. Thus we have roughly

$$\mathrm{SE}(\mathcal{Z}, \mathcal{S}) \sim 1 - a/\pi \tag{S11}$$

From this we see that in the limit where the cutoff is small and the simulation time is large, BE $\to 1$. On the other hand, for larger cutoff $\eta$ and shorter time scales $T$, the extinction boundary occurs at some fixed negative $\theta_1 \sim (\ln\eta)/T$ (with initial effects shifting $\ln\eta, T$ by small finite constants), and the difference in area will go as roughly $a(1-\theta_1^2/2)$, so we expect an effect something like

$$\mathrm{BE}(\mathcal{Z}, \mathcal{S}) = \mathrm{SE}(\mathcal{Z}, \mathcal{S})/\mathrm{LE}(\mathcal{Z}, \mathcal{S}) \sim 1 + a(\ln\eta)^2/(2t^2)\,. \tag{S12}$$

An example of this is shown in Figure S2. Thus, the perturbative analysis suggests that for reasonably short time scales and appropriate cutoffs, the linearized effect will lead to BE being positive (negative) when AE is negative (positive), as seen in many of the other examples in the paper. Nonlinear effects, however, will play an important role in multi-species systems with strong interactions, and the detailed relationship in general between BE and AE depends on the precise cutoff and time scale used to compute SE and appears to be relatively complicated for systems outside the perturbative regime.

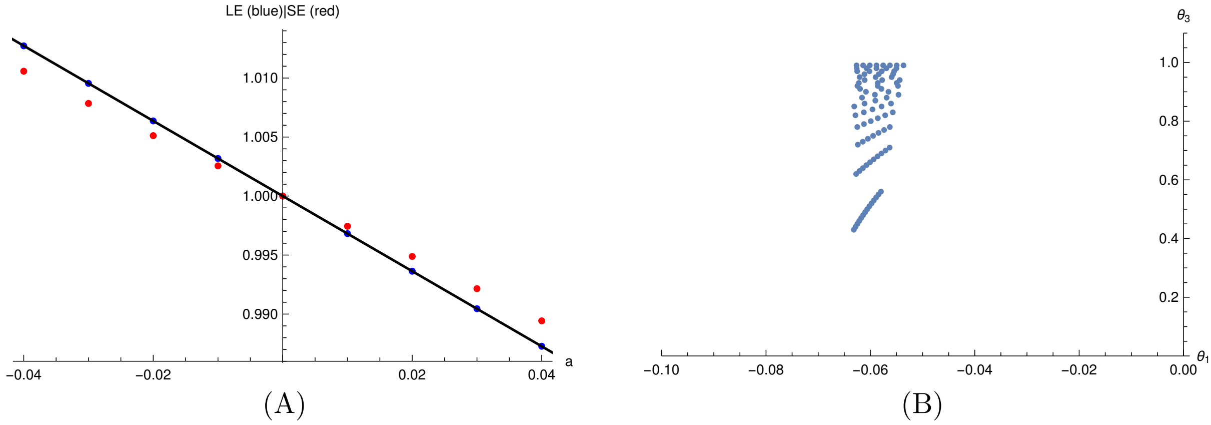


Supplementary Figure S2: **Long-term and short-term effects in the perturbative regime.** The effect of a third species on a pair of species (1, 2) with small interaction parameter $a \neq 0$ and other interactions vanishing in (S7). In Panel (A), black line is the perturbative theoretical prediction for LE (S8), and blue dots are exact analytic values of LE. Red dots are simulated effects SE, for extinction threshold $\eta = 0.0001$ and time $T = 100$, using 125,000 environmental conditions $\theta$ uniformly distributed on the unit sphere and initial conditions with all populations at 0.5. This example illustrates the general trend that LE $> 1$ ($< 1$) is correlated with BE $=$ SE/LE $< 1$ ($> 1$). Panel (B) depicts the set of initial conditions where the persistence of species (1, 2) differs in the presence or absence of the third species (3) over the finite time simulations, in the case $a = 0.01$, illustrating the shape expected from the theoretical analysis. Detailed pattern of points reflects sampling choice over sphere (grid points uniformly spaced in $\theta_3, \tan^{-1}(\theta_2/\theta_1)$); extinction boundary is localized near $\theta_1 \sim -0.06 \sim (\ln \eta)/T$ (up to finite shift of $\ln \eta, T$ from initial conditions), and expands in width roughly as $\theta_3$ as predicted.

#### S6 Supplementary Figures

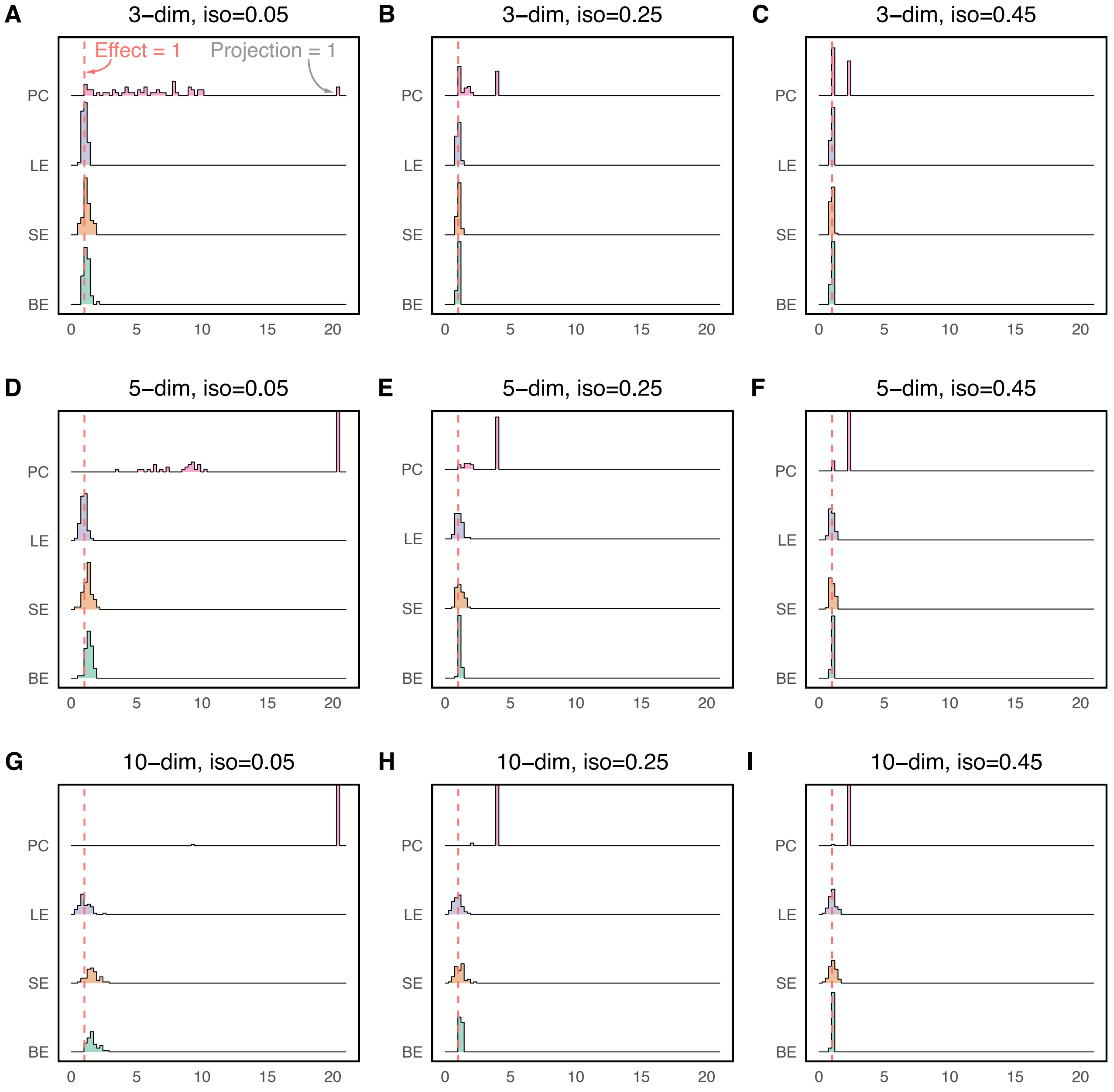


Supplementary Figure S3: **Distribution of system-level effects calculated on the fixed pair in Fig. 1D with different feasibility in isolation when embedded in 50 random systems in different dimensions where the interactions associated with other species in the systems are chosen randomly from a normal distribution with $\mu = 0$ and $\sigma = 0.25$.** PC means projection contribution; LE means long-term effects; SE means short-term effects; BE means buffering effects. In Panels **A, B, C**, the 50 random systems are 3-dimensional; in Panels **D, E, F**, the 50 random systems are 5-dimensional; in Panels **G, H, I**, the 50 random systems are 10-dimensional. In Panels **A, D, G**, the feasibility of the fixed pair in isolation (iso) is 0.05; in Panels **B, E, H**, the feasibility of the fixed pair in isolation is 0.25; in Panels **C, F, I**, the feasibility of the fixed pair in isolation is 0.45. For reference, the dashed line shows the value of one: the relative feasibility in isolation. The conclusions are summarized under Fig. S5 where the sampling distribution of the interactions associated with other species in the systems has a standard deviation of 1 (i.e., $\sigma = 1$).

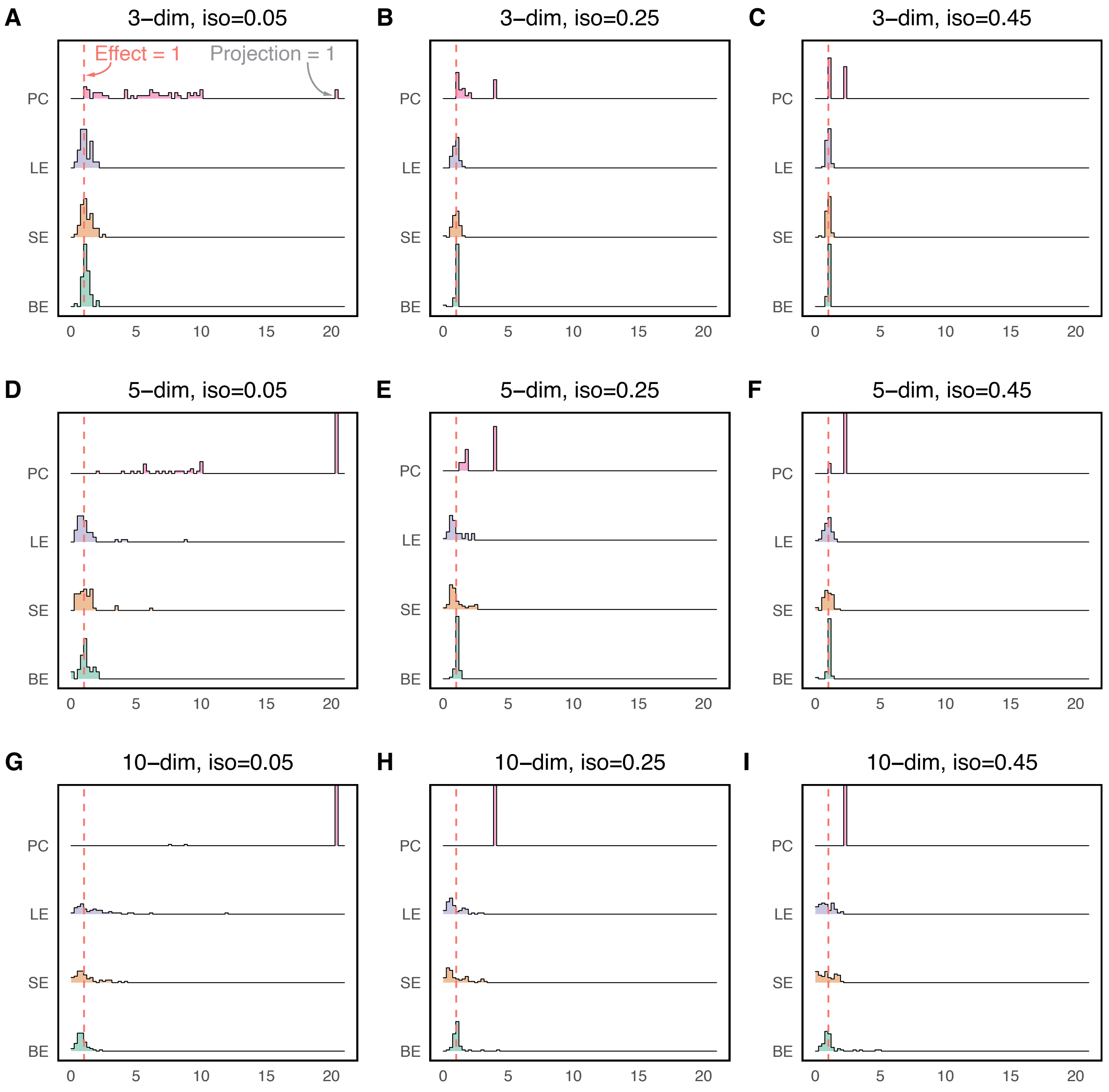


Supplementary Figure S4: **Distribution of system-level effects calculated on the fixed pair in Fig. 1D with different feasibility in isolation when embedded in 50 random systems in different dimensions where the interactions associated with other species in the systems are chosen randomly from a normal distribution with $\mu = 0$ and $\sigma = 0.5$ (see Fig. S3).** Here, the legend and layout of panels are the same as Fig. S3. The conclusions are summarized under Fig. S5 where the sampling distribution of the interactions associated with other species in the systems has a standard deviation of 1 (i.e., $\sigma = 1$).

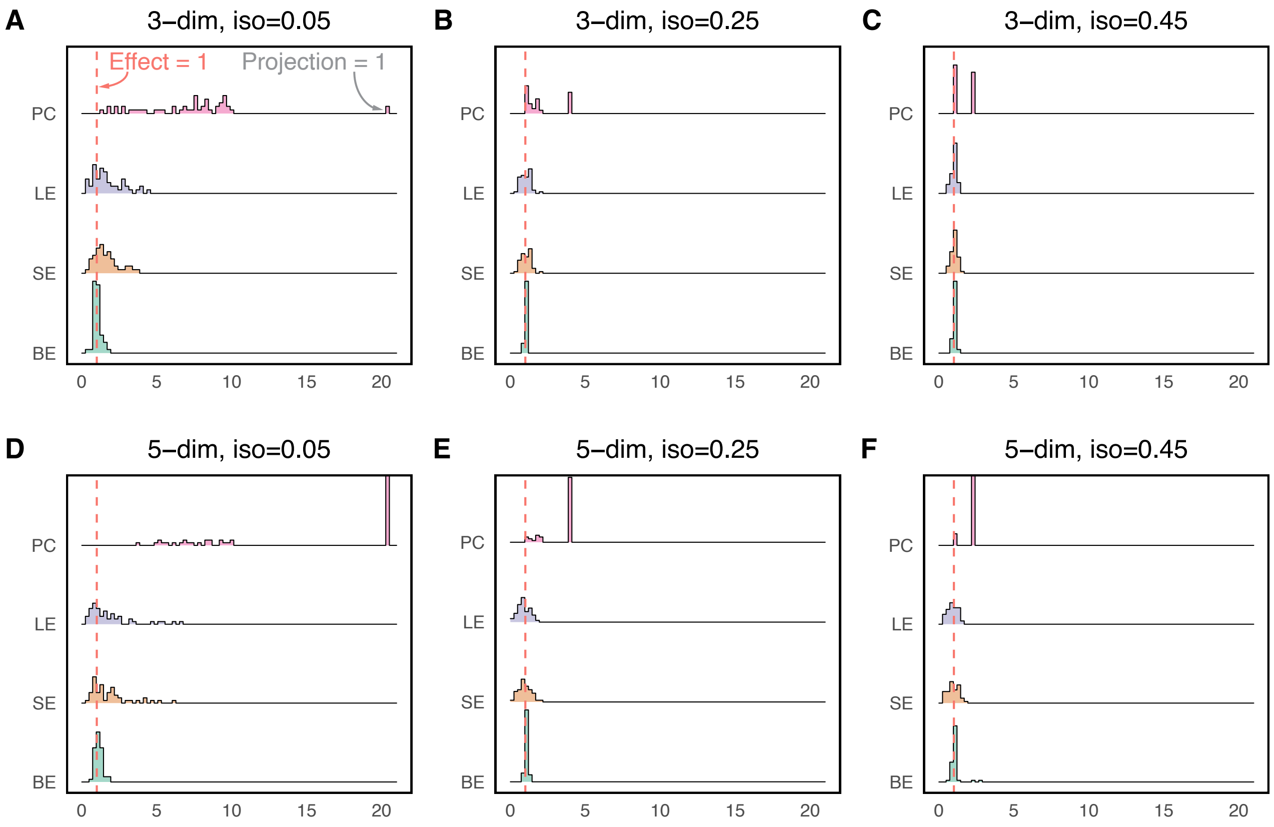


Supplementary Figure S5: **Distribution of system-level effects calculated on the fixed pair in Fig. 1D with different feasibility in isolation when embedded in 50 random systems in different dimensions where the interactions associated with other species in the systems are chosen randomly from a normal distribution with $\mu = 0$ and $\sigma = 1$ (see Figs. S3-S4).** Here, the legend and layout of panels are the same as Fig. S3. The following conclusions are based on Figs. S3-S5. The distribution of projection contribution (PC) clusters at the maximum (when projection = 1) as the dimension of systems (dim) increases. The range of PC (width of distribution) decreases as the feasibility of the pair in isolation (iso) increases. The distributions of long-term effects (LE), short-term effects (SE), and buffering effects (BE) tend to have larger widths as either the dimension of systems (dim) or the standard deviation of the sampling distribution of interactions ($\sigma$) increases, or as the feasibility of the pair in isolation (iso) decreases. According to statistical tests at the 0.05 level of significance, the distribution of long-term effects (LE) is centered at 1 and the mean of buffering effects (BE) is larger than 1 (except for one case in Panel **G** of Fig. S4).

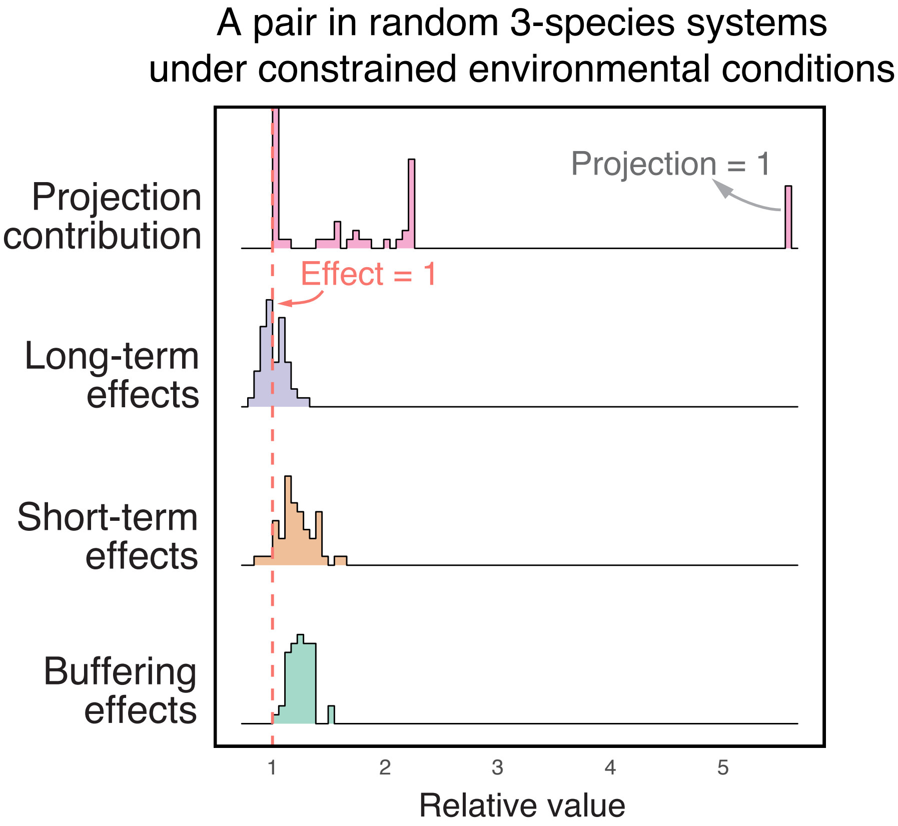


Supplementary Figure S6: **The distribution of system-level effects calculated on the fixed pair in Fig. 1D when embedded in 50 different 3-dimensional systems under constrained environmental conditions.** In the main text, we assume that the heterogeneous environments are completely unknown, so the effective growth rates $\theta$ of all species in the systems are uniformly distributed on the unit sphere (Fig. 1). Under environmental constraints, here we assume that the growth rate of individual species ranges between $[-0.9, 1]$. Each point in the distributions corresponds to the same pair in a different system. As a reference, the interaction matrix of the pair in isolation is $\begin{pmatrix} 1 & 0.22 \\ 0.22 & 1 \end{pmatrix}$.

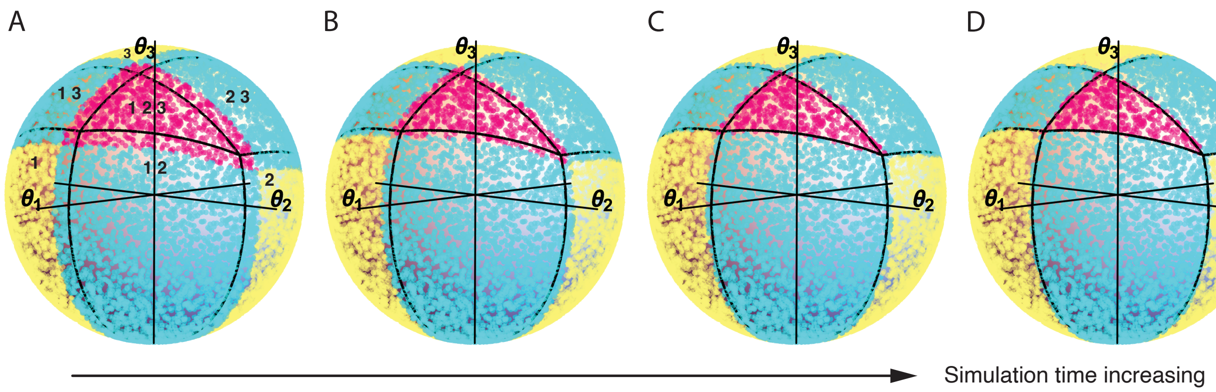


Supplementary Figure S7: **Resulting species compositions from 10,000 simulations over different times (see Fig. 1C).** The total time of simulations in Panels **A**, **B**, **C**, and **D** are 100, 200, 400, and 800, respectively. The step size is fixed as 0.01. The simulations are conducted by the Runge-Kutta method. The black lines correspond to the borders of the analytical feasibility regions in Fig. 1B. As time increases, the points on the sphere tend to cluster in the corresponding analytical feasibility regions. As a reference, the interaction matrix of the 3-species system is $\begin{pmatrix} 1 & 0.22 & 0.56 \\ 0.22 & 1 & 0.37 \\ 0.56 & 0.37 & 1 \end{pmatrix}$.

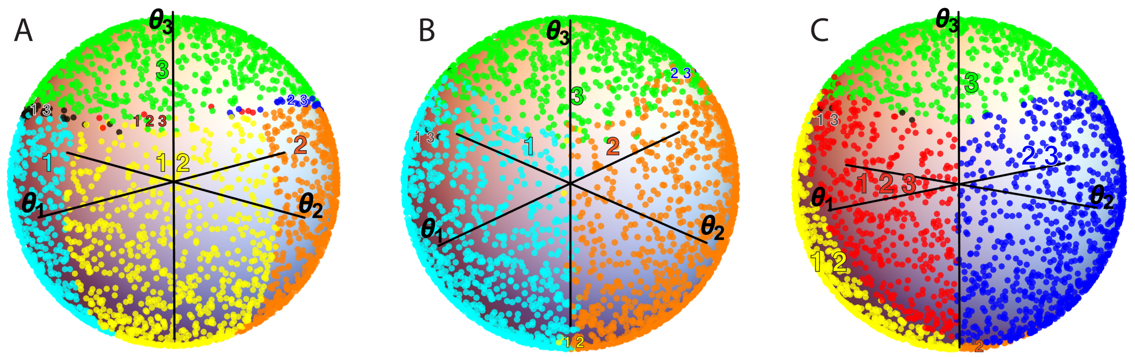


Supplementary Figure S8: **Resulting species compositions from 5,000 simulations with large interspecific interactions.** In Panel **A**, species 3 has strong harmful effects on species 1 and 2. In Panel **B**, all three species have strong harmful effects on each other. In Panel **C**, species have either strong harmful or strong beneficial effects on each other. In the numerical settings of all cases, the total time is 200, step size is 0.01 and extinction threshold is $10^{-6}$. The simulations are conducted by the Runge-Kutta method. As a reference, the interaction matrices in Panels **A, B, and C** are $\begin{pmatrix} 1 & 0.22 & 1.56 \\ 0.22 & 1 & 1.37 \\ 0.7 & 0.8 & 1 \end{pmatrix}$, $\begin{pmatrix} 1 & 1.22 & 1.56 \\ 1.22 & 1 & 1.37 \\ 1.7 & 1.8 & 1 \end{pmatrix}$, and $\begin{pmatrix} 1 & 1.22 & 1.56 \\ -1.22 & 1 & 1.37 \\ 1.7 & -1.8 & 1 \end{pmatrix}$, respectively. Also, the three systems here are not globally stable.

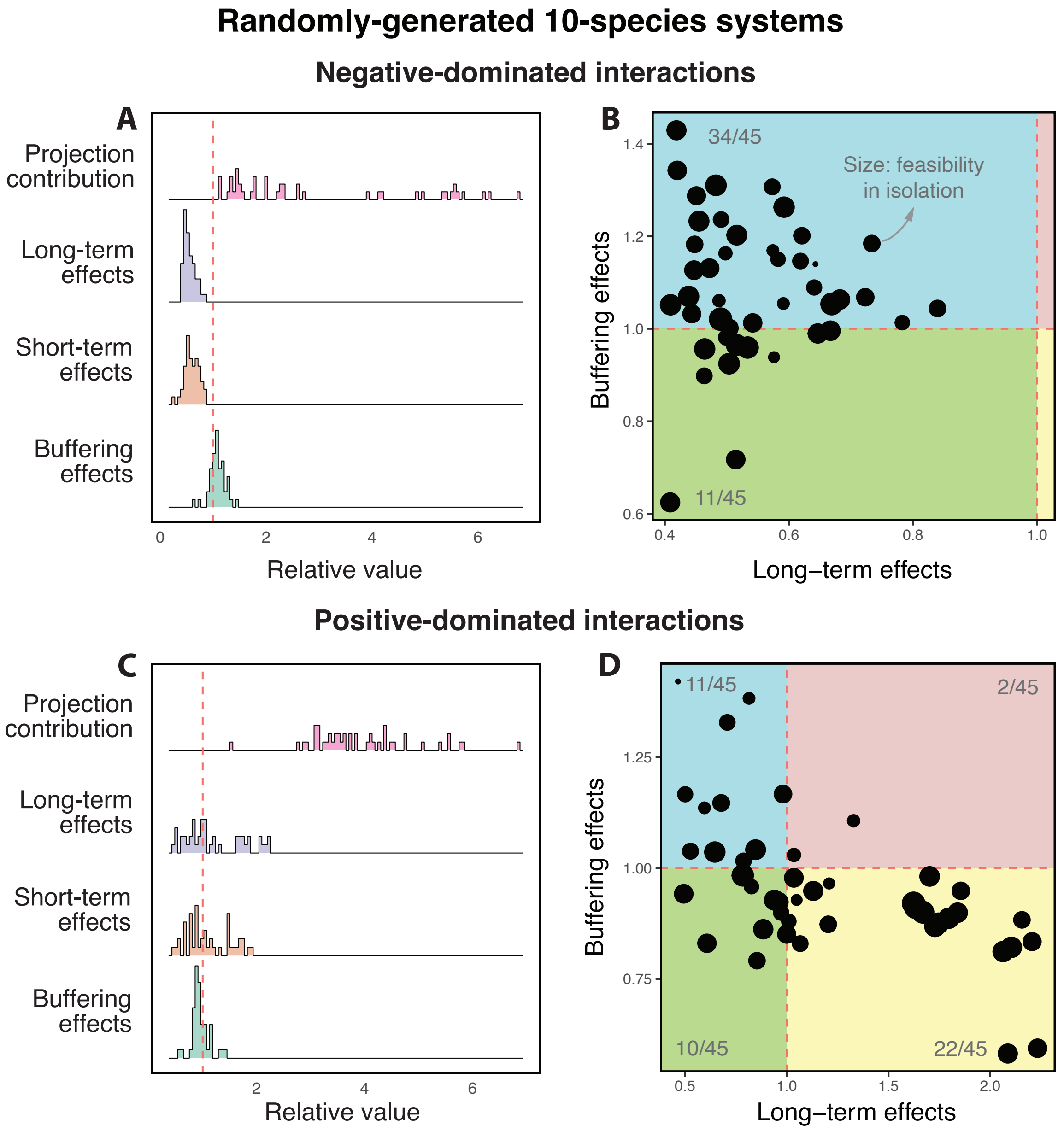


Supplementary Figure S9: **Cartographic representation of long-term and short-term differences among pairwise coexistence within the same 10-species system.** Panels **A** and **C** show the system-level effects on 45 pairs within a randomly-generated ten-species system characterized by negative-dominated and positive-dominated interactions, respectively (see text). Rows correspond to the projection distribution ($PC(\mathcal{Z}, \mathcal{S})$), long-term effects ($LE(\mathcal{Z}, \mathcal{S})$), short-term effects ($SE(\mathcal{Z}, \mathcal{S})$), and buffering effects ($BE(\mathcal{Z}, \mathcal{S})$), respectively. Note that each point in the distributions is a different pair within a system. For reference, the dashed line shows the value of 1. Recall that the x-axis corresponds to the change in probability of pairwise coexistence within the system. Panels **B** and **D** illustrate the cartographic representation for all corresponding pairs based on beneficial ($LE(\mathcal{Z}, \mathcal{S}) > 1$) or detrimental ($LE(\mathcal{Z}, \mathcal{S}) < 1$) long-term effects and beneficial ($BE(\mathcal{Z}, \mathcal{S}) < 1$) or detrimental ($BE(\mathcal{Z}, \mathcal{S}) < 1$) short-term effects. The size of points corresponds to the feasibility of pairs in isolation. The number of points in each region is annotated in gray.

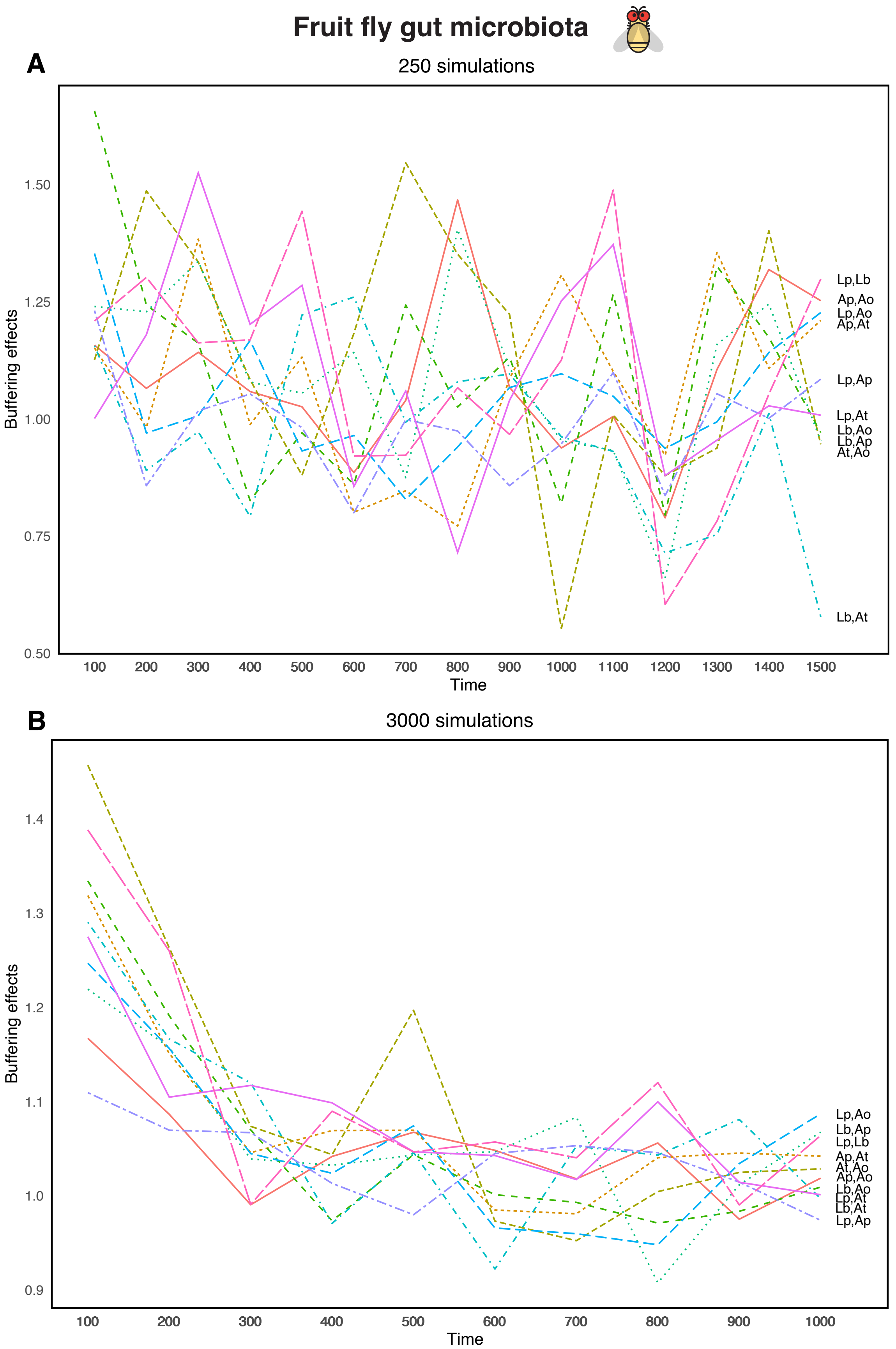


Supplementary Figure S10: **Buffering effects obtained by simulating dynamics over time.** As time increases, the distribution of buffering effects (y-axis) of the system on pairwise coexistence gradually shifts to a position centered at zero. The shift is faster and smoother as the number of simulations increases (Panel **A**: 250 simulations; Panel **B**: 3000 simulations). In the numerical settings of both cases, the size of time step (x-axis) is fixed as 0.01 and the extinction threshold is $10^{-6}$. Here, the interaction matrix of fruit-fly experiments is inferred as shown in Sec. S2.

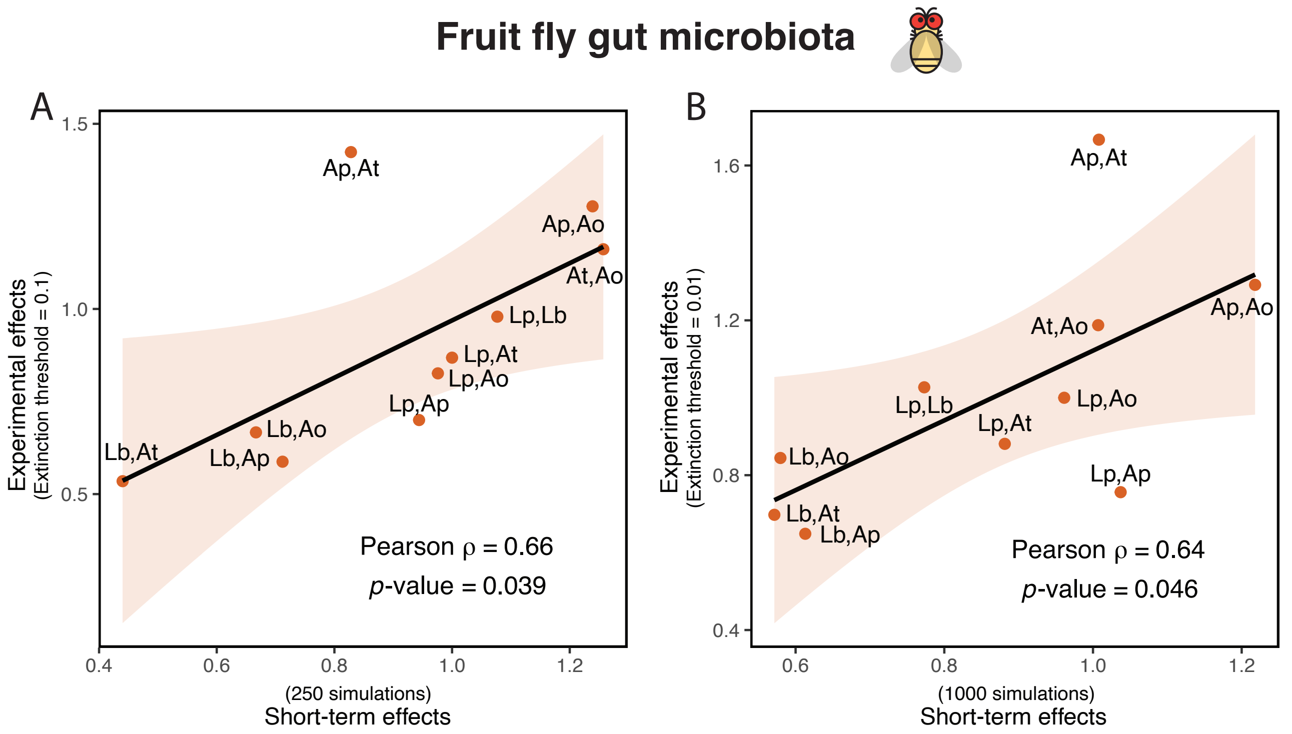


Supplementary Figure S11: **Correlation between experimental (short-term) effects and (theoretical) short-term effects.** When calculating the experimental effects (y-axis), we classify a species statistically extinct in a trial if its relative abundance was less than 1% (Panel **A**) and 10% (Panel **B**), respectively, across all replicates. The short-term effects (x-axis) are obtained by simulating the Lotka-Volterra dynamics Eq. (S1) using our inferred interaction matrix in Sec. S2 for 250 times (Panel **A**) and 1000 times (Panel **B**), respectively. In the numerical settings of both cases, the total time is 200 with step size 0.01 and the extinction threshold is $10^{-6}$. The simulations are conducted by the Runge-Kutta method. We found strong Pearson correlations (Panel **A**: $\rho = 0.66,\ p{-}\text{value} = 0.039$; Panel **B**: $\rho = 0.64,\ p - \text{value} = 0.046$) between the experimental effects and the short-term effects. Thus, with suitable parameters (e.g. number of simulations, extinction threshold, simulation runtime), the numerical simulations can successfully capture the information of pairwise coexistence within multispecies systems in the experimental settings.

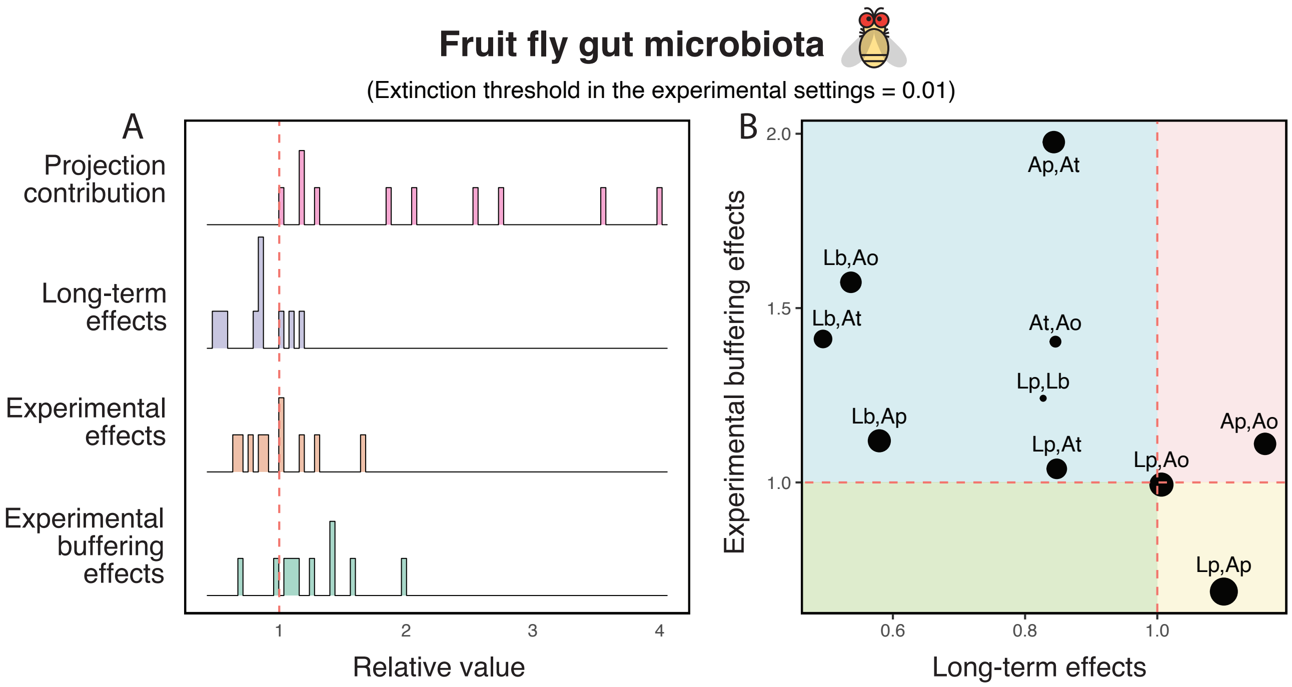


Supplementary Figure S12: **Comparing system-level effects among pairs within a five-species *in vivo* experimental system.** Compared to Fig. 4 in the main text, here we classify a species statistically extinct in a trial if its relative abundance was less than 10% across all replicates (1% in Fig. 4).
